# Cytomegalovirus Infection in Human Cortical Organoids Induces Unique Developmental and Viral Substates

**DOI:** 10.64898/2026.11.04.764258

**Authors:** Daniel A. Laws, Kaitlyn J. Partridge, Benjamin S. O’Brien, Chien-Wei Lin, Kenichiro Taniguchi, Allison D. Ebert, Scott S. Terhune

## Abstract

Human Cytomegalovirus (HCMV) is a ubiquitous β-herpesvirus with exposure resulting in a life-long infection. Severe HCMV disease can occur in conditions of insufficient immunity, and this includes congenital infection. Infection of the developing brain can cause severe birth defects including microcephaly and sensorineural hearing loss. Also, increasing evidence suggests the contribution of HCMV infection in adult CNS diseases. Studying HCMV replication and its impact on CNS is restricted by host tropism and anatomy. In these studies, we define the impact of infection on healthy iPSC-derived cortical organoids using single cell RNA sequencing. We uncovered 16 cell type clusters in uninfected tissues using the Human Neural Organoid Cell Atlas. Following infection by HCMV expressing GFP, several clusters were altered in both GFP-positive and GFP-negative populations, including the likely transition of the Glutamatergic/GABAergic Neuron cluster into a CMV-Infected Cell cluster. Expressions of viral genes varied between cells and cell types, requiring us to introduce a TC (temporal class) indexing score to account for varying stages of infection. This analysis in conjunction with machine learning approaches revealed cells that are likely producing de novo HCMV along with several viral sub-states. Several viral sub-states had significant enrichments of genes involved in epithelial-to-mesenchymal (EMT) transition, including expression of EpCAM. We independently verified HCMV-induction of EpCAM protein expression in cortical organoids and glutamatergic neurons. In summary, our studies reveal unique features of HCMV infection of cortical organoid tissues, including varying viral sub-states and tropisms, and introduced new approaches to analyze these multi-dimensional complex data sets.

**SUMMARY:** Congenital HCMV infection can lead to significant neuropathological consequences. Here we analyzed human stem cell derived cortical organoids by single cell RNA sequencing and subsequent machine learning algorithms to identify and classify viral progression and neural cell responses to HCMV infection. These data help elucidate infection trajectory and the transcriptional landscape across a complex human neural model system.

## INTRODUCTION

Human cytomegalovirus (HCMV) is a ubiquitous β-herpesvirus with a global seroprevalence ranging between 70-90% (1). While HCMV infection in immunocompetent adults is usually self-limiting, primary infection, infection by a second variant, or latent reactivation in immunocompromised individuals can result in severe complications. During pregnancy, primary HCMV infection or re-infection is the leading cause of nonheritable birth defects including sensorineural hearing loss, microcephaly, neurodevelopmental delays, and seizures (2). Imaging of fetal brains reveals loss of the cortical intermediate zone, focal necrosis, hemorrhage, and periventricular calcification, all contributing to structural brain abnormalities at birth (3). Current antiviral pharmaceuticals targeting HCMV, such as ganciclovir and valganciclovir, are limited due to significant side effects and are not indicated for use during pregnancy due to their teratogenicity. However, use of the antiviral valacyclovir during pregnancy has emerged as an approach to reduce vertical transmission (4,5). These therapeutics do lack specificity for latent HCMV reservoirs, permitting reactivation in immunocompromised patients. Finally, attempts to develop a vaccine against HCMV continue to be unsuccessful including a recent stopped Phase 3 CMVictory trial (NCT05085366) of an mRNA vaccine (6). Considering the burden of complications associated with infection, HCMV remains a high-level priority for further study.

HCMV is species-specific, requiring human-derived samples to study infection and pathogenesis. For studies on neuropathogenesis, human induced pluripotent stem cells (iPSCs) have become a valuable model system as they can be differentiated into individual neural cells and 3-dimensional organoid structures to model brain development (7). Studies using neural progenitor cells have discovered critical roles for HCMV proteins in altering cell proliferation (8), migration (9), and altered cell fates (10). Recently, we demonstrated that HCMV infects and replicates in terminally differentiated glutamatergic neurons (11,12). Infection resulted in notable neurite retraction, formation of multinucleated syncytia, and loss of neuronal action potentials. Human iPSCs have also been used to generate neural organoids consisting of cells in varying developmental states. Single cell RNA sequencing (scRNA-seq) has shown that iPSC-derived cortical organoids generate the specific cell types found in the developing human brain, including neural progenitor cells, excitatory and inhibitory neuron populations, and various glial cell populations (13). We and others have used iPSC-derived cortical organoids to elucidate several pathways dysregulated following HCMV infection (14–22). These studies used a variety of different iPSC lines, methods of organoid generation, infections at different days of development from 15 to 75 days, and HCMV TB40 or Merlin strains with varying conditions of infection. Emerging themes in developmental disruption include destabilization of tissue architecture, dysregulation of neurodevelopmental programs, and secondary injury.

Following infection of day 30 organoids by an inoculum determined by tissue weight, we have observed downregulation of key neurodevelopmental pathways in both an HCMV IE1-dependent and -independent manner by 14 dpi, and significant transcriptional changes in cell populations with little to no viral gene expression. Transcriptionally downregulated neurodevelopmental genes observed in several studies in either organoids or neuronal cells by multiple groups include HES1 (19,23–25), NESTIN (26–28), SOX2 (19,26,27,29) and OLIG1 (19,26), which also coincides with significant disruption in structural organization and calcium signaling (19,21,26,30). In addition to developmental genes, infection stimulated numerous interferon-regulated genes including MX1, OAS1, and ISG15 (19,31,32).

Lytic HCMV replication in monolayer fibroblast cells occurs in a coordinated series of steps that have been experimentally defined *in vitro* (Reviewed in Goodrum *et al.* 2022 (33)). Classification of each step is based on experimental perturbation and influenced by the method of detection and timepoints (34,35). Genes expressed in the Immediate Early (IE) cluster are resistant to cycloheximide-mediated inhibition of translation. The Late (L) gene cluster is defined by expressions sensitive to viral genome synthesis inhibitors. The Early (E) gene clusters exhibit expressions sensitive to cycloheximide but insensitive to replication inhibitors, and there are several additional clusters that were defined based on intermediate and kinetic phenotypes. Moreover, viral gene classifications vary based on methods of detection (e.g., RNA microarray, RNA-seq) (34,35). For example, a gene defined as an Early gene by microarray might be reclassified as an Immediate Early by RNA-seq due to using a more sensitive analysis method. The use of RNA-seq identified additional clusters, referred to as Temporal Classes, with each gene clustered based on the pattern of expression over time. However, the expression levels between different genes in the same cluster cannot be compared as levels over time are set relative to the highest point for each gene. These studies have been completed using synchronized monolayer cultures of a single cell type, which leaves uncertainty as to how viral classification could be impacted by an asynchronous infection in a three-dimensional tissue structure containing multiple distinct cell types.

Here, we aimed to define the impact of HCMV infection in an iPSC-derived cortical organoid model using scRNA-seq and spatial transcriptomics. We evaluate the asynchronous infection and how it impacts cell type developmental states using standard approaches in scRNA-seq data analyses. Due to the multi-dimensional nature of the data (e.g. cell developmental states, stages of infection, three-dimensional tissue, etc.), we also apply several machine learning approaches. We have uncovered varying viral-sub-states within the infected tissues which have broad implication relating to HCMV disease.

## RESULTS

### HCMV infection alters cellular identities in human cortical organoids

HCMV infection damages the developing brain with increasing evidence of a negative impact on adult brains (36,37). To define how infection alters developmental states, we infected healthy control iPSC-derived cortical organoids. Organoid cultures were differentiated using a cerebral organoid kit based on an established protocol (38), which was used in our previous studies (16,18,19,21). At 30 days post differentiation, we mock-treated or infected tissues using BAC-derived HCMV strain TB40 containing the eGFP gene at 500 IU/µg of tissue (**Fig 1A**)(16,18,19). Tissues were dissociated at 14 days post-infection (dpi) into single cell suspensions and analyzed by scRNA-seq. To uncover HCMV-induced developmental states, we analyzed mock and HCMV-infected GFP-negative and GFP-positive scRNA-seq datasets independently to avoid masking any potential unique states. The resulting sequencing data were aligned to the human reference genome g19 and sequences from HCMV TB40/E (39). We performed filtering and normalization in Seurat (v5.4), and we used the Louvain algorithm to identify transcriptional clusters. We then colored cells on the UMAP projection by cluster identity for all conditions using Scanpy (v1.11.5).

**Fig 1.**
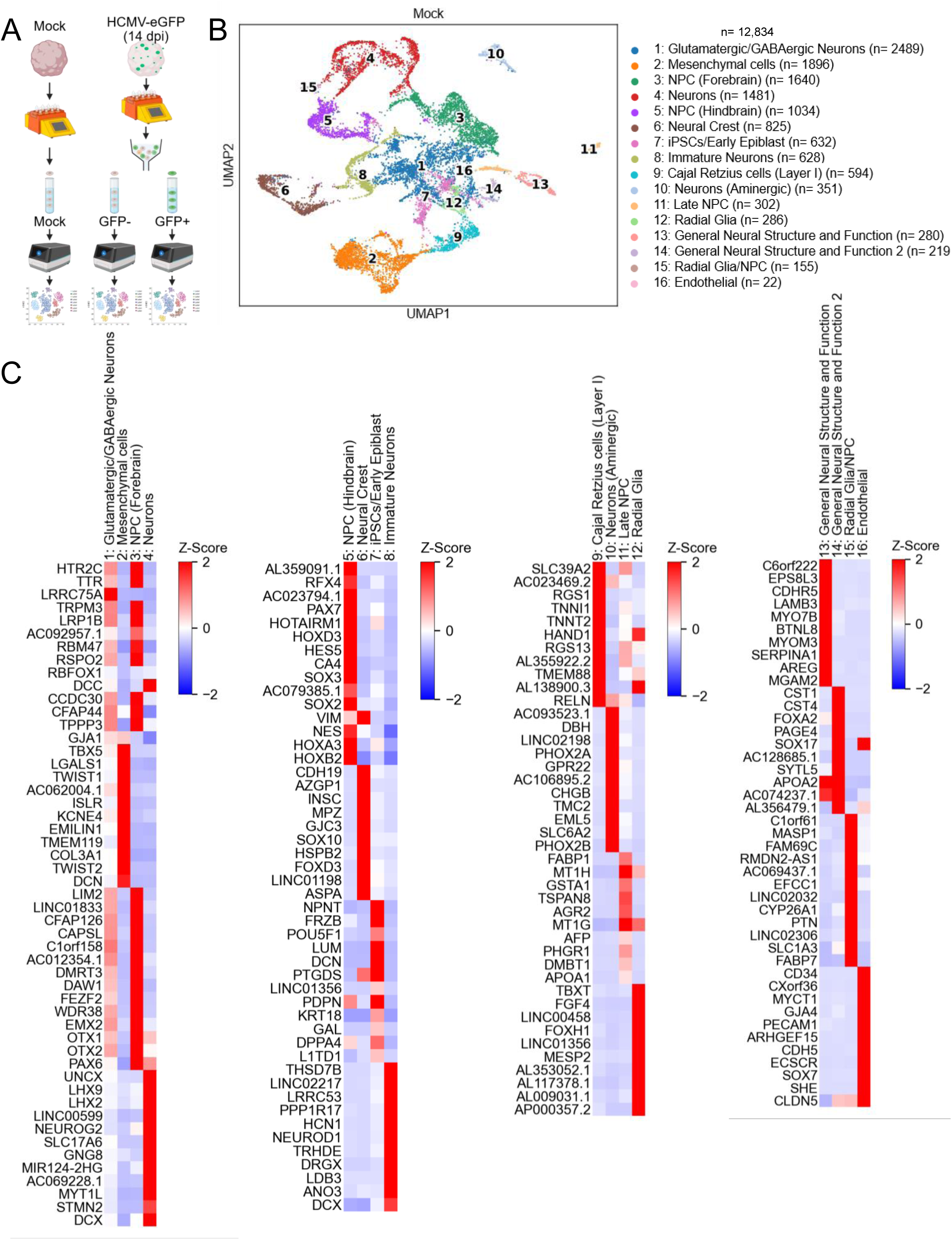
Cell type clusters and gene marker heatmaps of scRNA sequenced cortical organoids. (A) Thirty day old iPSC-derived cortical organoids were mock-treated or infected using BAC-derived HCMV TB40 strain encoding the green fluorescent protein (GFP). At 14 dpi, organoids were dissociated with infected organoids sorted into GFP-positive and GFP-negative expressing cells using by flow cytometry. Samples subsequently underwent single cell RNA sequencing. (B) UMAP plot of HCMV uninfected “Mock” iPSC-derived cortical organoids clustered by cell type. (C) Heatmap showing host markers used to define and annotate UMAP clusters.

For mock-treated organoids, we identified 16 cell type clusters (**Fig 1B**). Cluster identities were determined using the Human Neural Organoid Cell Atlas (HNOCA) developed by He *et al.* (40) and canonical marker genes to identify specific clusters (40,41). If there was no clear signature, we next identified the 20 differentially expressed (DE) genes and submitted the list to AnnDictionary (42), querying with Claude Haiku 4.5 (43) to provide putative cluster annotation. Where possible, specific names of the cell type clusters were derived directly from the noted references. The most abundant clusters in Mock-treated cortical organoids include Glutamatergic/GABAergic Neurons (gene markers HTR2C, LRP1B), NPC (Forebrain) (FEZF2, EMX2, OTX2, PAX6), Mesenchymal cells (TBX5, TWIST1, TWIST2), NPC (Hindbrain) (HOXD3, HOXB1, HES5, HOXB2) and Neurons (DCX, NEUROG2, STMN2, LHX9) (**Figs 1B and S1B Fig**). Other cluster identities include IPSCs/Early Epiblast, Immature Neurons, Cajal Retzius cells, Neural Crest, Neuron (Aminergic), Late NPC, Radial Glia, and Radial Glia/NPC along with two heterogeneous clusters which we have termed General Neural Structure and Function 1 and 2. A small number of cells fell into an Endothelial cluster in all conditions despite organoids transitioning via a neuroectodermal stage. We plotted the average expression of the top marker genes across all cells in a given cluster in heatmaps to compare unique expressions between clusters (**Fig 1C**). To evaluate specificity of each set of marker genes (40–42,44) for their respective annotated cluster, we calculated a module scored expression for each gene set using built in module scoring functions from the Scanpy library. Module scores between clusters are compared using a violin plot with scores significantly elevated in each respective cluster relative to the others (**S1A Fig**).

Infection of cortical organoids using TB40-eGFP resulted in a fraction of cells exhibiting fluorescence (**Fig 2A**). As done previously (19), we sorted cells based on GFP fluorescence intensity into GFP-positive cells and GFP-negative cells, with gates set using uninfected mock cells. We refer to this latter population as GFP-negative based on the lack of detectable fluorescence. We completed scRNA-seq on both GFP-positive cells and GFP-negative cells with resulting data analyzed separately. We identified 10 cell type clusters (**Fig 2B**) in the GFP-negative group. We plotted the average expression of the top marker genes across all cells in each cluster with heatmaps to compare unique expressions between clusters (**S2A Fig**) and calculated a module scored expression for each gene set (**S2B Fig**). Using the same approach as above, we defined cluster identities for the majority of cells belonging to Mesenchymal, Neurons 1, Neurons 2, Ependymal Cells, and NPC clusters (**Figs 2B** and **S2C Fig**). Other clusters include Neural Crest, Neurons (Aminergic), Oligodendrocytes, and Endoderm/Mesoderm Lineage.

**Fig 2.**
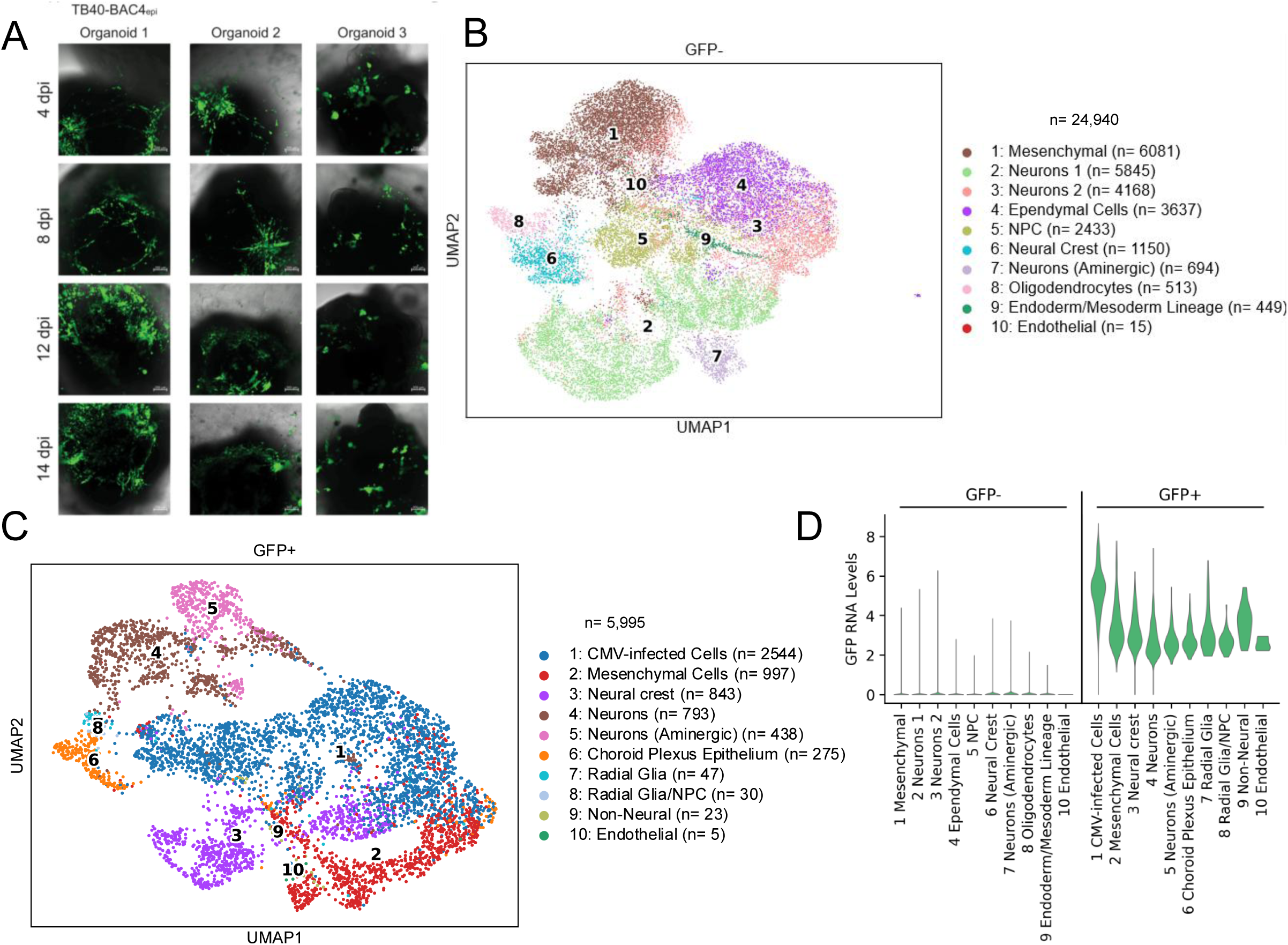
scRNA-seq of HCMV infected hiPSC-derived cortical organoids. (A) Immunofluorescence images showing GFP expression in HCMV infected cells across 3 independent organoids. (B) UMAP plot of the GFP-negative subset of the HCMV-infected neural organoid, clustered by cell type. (C) UMAP plot of the GFP-positive subset of the HCMV-infected neural organoid, clustered by cell type. (D) Violin plot confirming GFP transcript levels are largely absent in the GFP-negative subset, and present in the GFP-positive subset.

We recovered approximately 20% of the cells as GFP-positive from HCMV-infected organoids (**Fig 2**). Using the clustering methods described above, we clustered GFP-positive cells into 10 groups (**Fig 2C**). Clusters were defined based on unique gene sets (**S3A Fig**) and modular scores (**S3B Fig**), and resultant annotations included Mesenchymal Cells, Neural Crest, Neurons, Neurons (Aminergic), Choroid Plexus Epithelium, Radial Glia, Radial Glia/NPC, and Non-Neural cluster. The largest cluster, which we annotated “CMV-infected cells”, expressed predominantly viral transcripts, with relatively few host transcripts thereby limiting our ability to identify the cell types of origin (**S3C Fig**). It is reasonable to conclude that these cells have lost their inherent identity due to infection. This CMV-infected cells cluster represented approximately 42% of the cells exhibiting GFP fluorescence (**S2C Fig**). The population is likely capable of producing virus, as the top 10 marker transcripts are classified as Early-Late and Late genes (UL84, UL53, UL83, UL76, UL52, UL74, UL22A, UL146, UL139) **(S3A Fig**) (35,45). Finally, to validate our separation of cells by GFP fluorescence, we compared GFP RNA levels between all clusters (**S2D Fig**). We detected GFP expression in the GFP-positive groups with the highest levels of expression in the CMV-infected cells cluster. In contrast, little to no GFP RNA was detected in the GFP-negative group, as seen in our previous bulk RNA-seq studies (19).

### Spatial transcriptomics of HCMV-infected 30 day organoids

We utilized spatial transcriptomics to map potential cellular responses to HCMV infection with the 10x Genomics Xenium *in situ* imaging platform. We infected and mock-treated 30 day old organoids as described above and embedded multiple organoids per block at 14 dpi. We cut 10 µm sequential sections for *in situ* hybridization, using sections 30 µm apart for spatial staining with two sections in between for GFP staining and quality control. We used confocal microscopy to image mock and HCMV-infected cortical organoids to confirm that HCMV-infected cells exhibit GFP positivity that is not seen in mock-treated samples (**Fig 3A**). Targeted spatial profiling was conducted across biological replicates using the Xenium Human Brain Gene Expression Panel. The panel consists of 266 gene targets including 89 related to neuronal development, 68 glioblastoma-related, 30 astrocytes or microglia-related, 26 vascular-related, and 35 oligodendrocyte-related.

**Fig 3.**
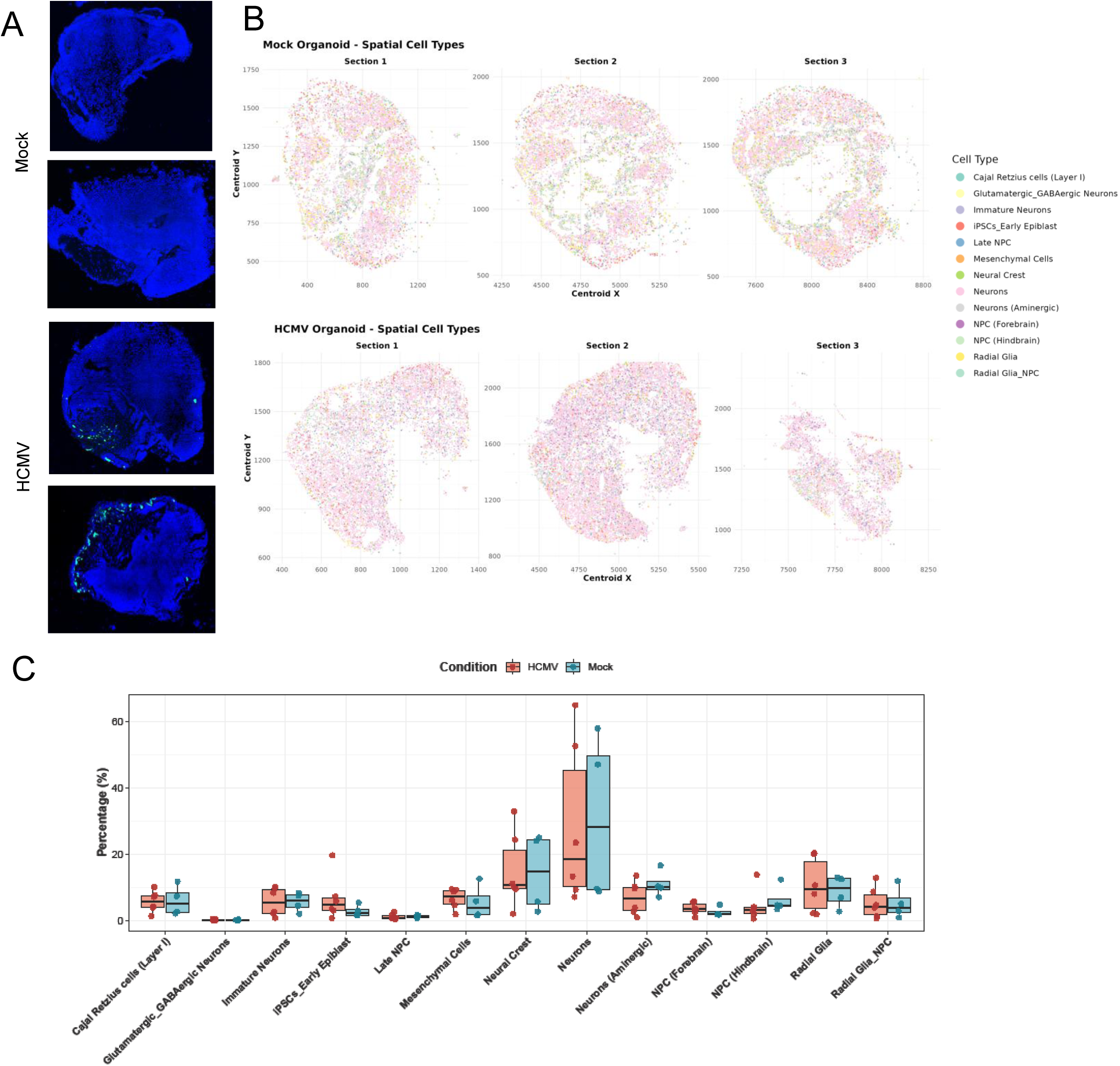
Spatial transcriptomics of mock and HCMV-infected cortical organoids. (A) Confocal microscopy images of the iPSC-derived cortical organoids used for spatial transcriptomics. HCMV-infected cells exhibit GFP fluorescence with two representative images shown. (B) Spatial UMAP projection of Mock and HCMV-infected cortical organoids, colored by cell type. (C) Box plot comparing cell type composition between all mock and all HCMV-infected cortical organoids.

The resulting spatial transcriptomics data were combined using the Robust Cell Type Decomposition (RCTD) algorithms (46), allowing for integration of our scRNA-seq dataset and spatial transcriptomic data, as well as mapping of cell-type composition across serial tissue sections. The successful deconvolution and confidence mapping of scRNA-seq reference cell types across our Xenium tissue sections suggest that our scRNA-seq reference reflects the cell populations present *in situ.* Additionally, our cortical organoid culture is consistent in producing organoids of similar cell type composition. However, although achieving high mapping confidence scores, there persists some inherent circularity. The organoids generate the diversity of lineage subsets, ranging from early epiblast to aminergic neurons (**Fig 3B**). We then plotted cell type composition of HCMV-infected and mock organoids used for spatial analysis to compare differences in overall cell type make-up (**Fig 3C**). We found no significant differences in cell type composition of HCMV-infected versus mock-treated organoids. We suspect that the limited number of genes in the array contributes to a lack of gene signature resolution necessary to define HCMV-induced developmental changes. In summary, spatial transcriptomic profiling of mock and HCMV-infected brain organoids validates that cellular identities defined by *in situ* hybridization of tissue sections closely correspond to cell populations identified within the scRNA-seq analysis.

### HCMV infection disrupts cell type states during development

A limitation of separately analyzing data sets is in identifying the cell-types of origin for HCMV-infected organoids (**Fig 2**) compared to control (**Fig 1**). Optimal Transport (OT) is a mathematical framework used to map probability distributions of different time point datasets onto one another while minimizing a transport cost function. (47). Multi-Omics Single-Cell Optimal Transport (MOSCOT) (v0.5.0) (48) is an adaptation of OT that can be used to map scRNA-seq datasets onto one another to transfer cluster annotations between datasets. Utilizing MOSCOT’s TemporalProblem class based on Schiebinger *et al.*, 2019 (38), we sought to model the transcriptional trajectories of mock cell types as they transition to GFP-negative (**Fig 4A**) and GFP-positive (**Fig 4B**) cell types following infection. In other words, scRNA-seq captures gene identities and expression levels to define cell states, and we used MOSCOT to compare how the patterns of different cell states under mock conditions moved or transitioned to those in infection-related conditions. In MOSCOT, we employed the TemporalProblem class (49) to facilitate the modeling of infection as a temporal perturbation to evaluate the probabilistic transcriptional mapping of mock subset cell type clusters onto the other subset cell type clusters. For example, a probability score approaching 1.0 means high biological certainty that a later cell state is a descendent of an earlier cell state when mapping development from a time course of scRNA-seq datasets.

**Fig 4.**
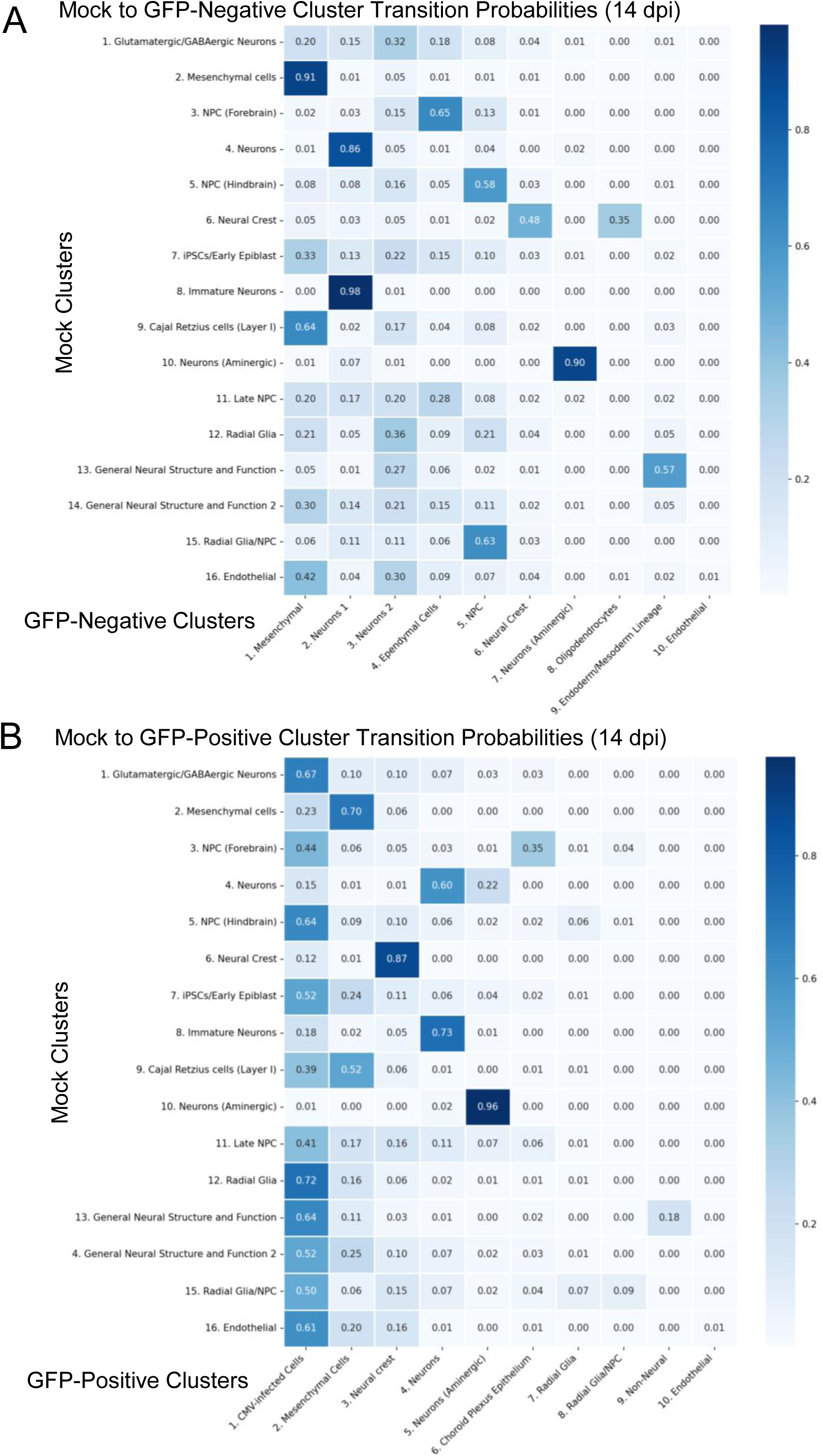
MOSCOT analysis of cell type transition probability following HCMV infection. Heatmap showing the cluster transition probability matrix from Multi-Omics Single-Cell Optimal Transport (MOSCOT) derived from solution of the TemporalProblem. Each row represents cell type clusters derived from Mock neural organoids, and each column represents cell type clusters derived from GFP positive subset (A) or GFP negative subset (B). Cells of the heatmap and their associated scores represent the proportion of aggregated probability mass of cells within a Mock cluster transitioning to cells within a GFP positive cluster. More generally, each cell of the heatmap corresponds to the probability that cells in a given GFP positive cluster transcriptionally originated from the cells of a particular Mock cluster following HCMV infection.

We plotted the probability matrix describing lineage fate of mock clusters moving to GFP-negative (**Fig 4A**) and GFP-positive clusters (**Fig 4B**). The transition probability is roughly proportional to the probability of mock cell type clusters transitioning to clusters following infection in our analysis. When reviewing Mock to GFP-negative probability scores (**Figs 4A and S4A Fig**), Mock Mesenchymal cells (Cluster 2) and Neurons (Aminergic) (Cluster 10) have a high probability of progressing to GFP-negative Mesenchymal (Cluster 1) and Neurons (Aminergic) (Cluster 7) at 0.91 and 0.90, respectively. For GFP-negative Neurons 1 (Cluster 2), our analysis suggests that cells in this cluster originate from a mix of Mock Neurons (Cluster 4) and Immature Neurons (Cluster 8) at probability scores of 0.86 and 0.98, respectively. In contrast, GFP-negative Neurons 2 (Cluster 3) and Ependymal Cells (Cluster 4) match with low probabilities to several Mock clusters, suggesting infection-mediated state changes despite low GFP expression levels (**Fig 2D**). Finally, we asked whether these GFP-negative cells were less susceptible to infection due to a lack of viral entry receptors (50). Analyses of gene expression for known receptors for HCMV trimeric, pentameric, and gB glycoprotein complexes showed mRNA expression of one or more entry receptors in each cluster (**S5 Fig**), likely ruling out this possibility.

For GFP-positive cells (**Fig 4B and S4B Fig**), Mock Neural Crest (Cluster 6) and Neurons (Aminergic) (Cluster 10) show high probabilities of progressing to GFP-positive Neural Crest (Cluster 3) and Neurons (Aminergic) Cluster 5 at 0.87 and 0.96, respectively. Cells in the GFP-positive Neurons (Cluster 4) appear to originate from Mock Neurons (Cluster 4) and Immature Neurons (Cluster 8) at 0.60 and 0.73, respectively, while cells in the Mesenchymal Cells (Cluster 2) originate from Mock Mesenchymal cells (Cluster 2) and Cajal Retzius cells (Cluster 9) at probabilities of 0.70 and 0.52, respectively. In contrast, analysis of GFP-positive CMV-infected Cells (Cluster 1) implicates multiple origins from multiple cell states found in Mock including Glutamatergic/GABAergic Neurons (Cluster 1), NPC (Hindbrain) (Cluster 5), and Radia Glia (Cluster 12). Interestingly, Mock Neural Crest (Cluster 6) and Neurons (Aminergic) (Cluster 10) have low probabilities of contributing to this GFP-positive CMV-infected Cells (Cluster 1) at 0.12 and 0.10, respectively. One limitation of projecting CMV-infected Cells (Cluster 1) back onto other states using MOSCOT, is that the high number of viral transcripts relative to cellular transcripts in these cells may bias the OT framework toward suggesting a high number of possible origins for these ambiguous cells. For this reason, it is important to consider the full picture of cell types that are conserved and absent between mock and GFP-positive/-negative organoids. In summary, this analysis suggests several cell states are conserved following HCMV infection (e.g., Aminergic Neurons, Neural Crest) whereas others are altered (e.g., Glutamatergic/GABAergic Neurons, Radial Glia) with the possibility of new states emerging, other than CMV-infected Cells cluster (e.g., Ependymal, Choroid Plexus Epithelium) (**S4C Fig**).

### Gradient expression of HCMV viral gene transcripts across different cell clusters

The 235 kbp HCMV genome can express hundreds of viral RNAs occurring in a temporal pattern during lytic infection including Immediate Early, Early, Early-Late and Late genes. This classification was further refined using bulk RNA-seq data following synchronized infections of fibroblasts into Temporal Classes (TC) 1 to 7 by Rozman *et al.* (45). Used in our analysis, TC1 (Immediate Early) consists of 10 viral genes, TC2 (Early) is 31 genes, TC3 (Early) is 38 genes, TC4 (Early-Late) is 9 genes, TC5 (Late, replication-independent) is 7 genes, TC6 (Late) is 7 genes, and TC7 (Late) is 22 genes (**S6 Fig**) (45). To understand HCMV gene expression patterns in the cortical organoids, we analyzed viral mRNA expressions by first quantifying total UMI count per cell in the GFP-positive population (**Fig 5A**). The highest total expression overlapped with the CMV-infected Cells (Cluster 1) followed by a fraction of cells in the Mesenchymal Cells (Cluster 2) and Neural crest (Cluster 3) with remaining clusters showing relatively low total UMI counts.

**Fig 5.**
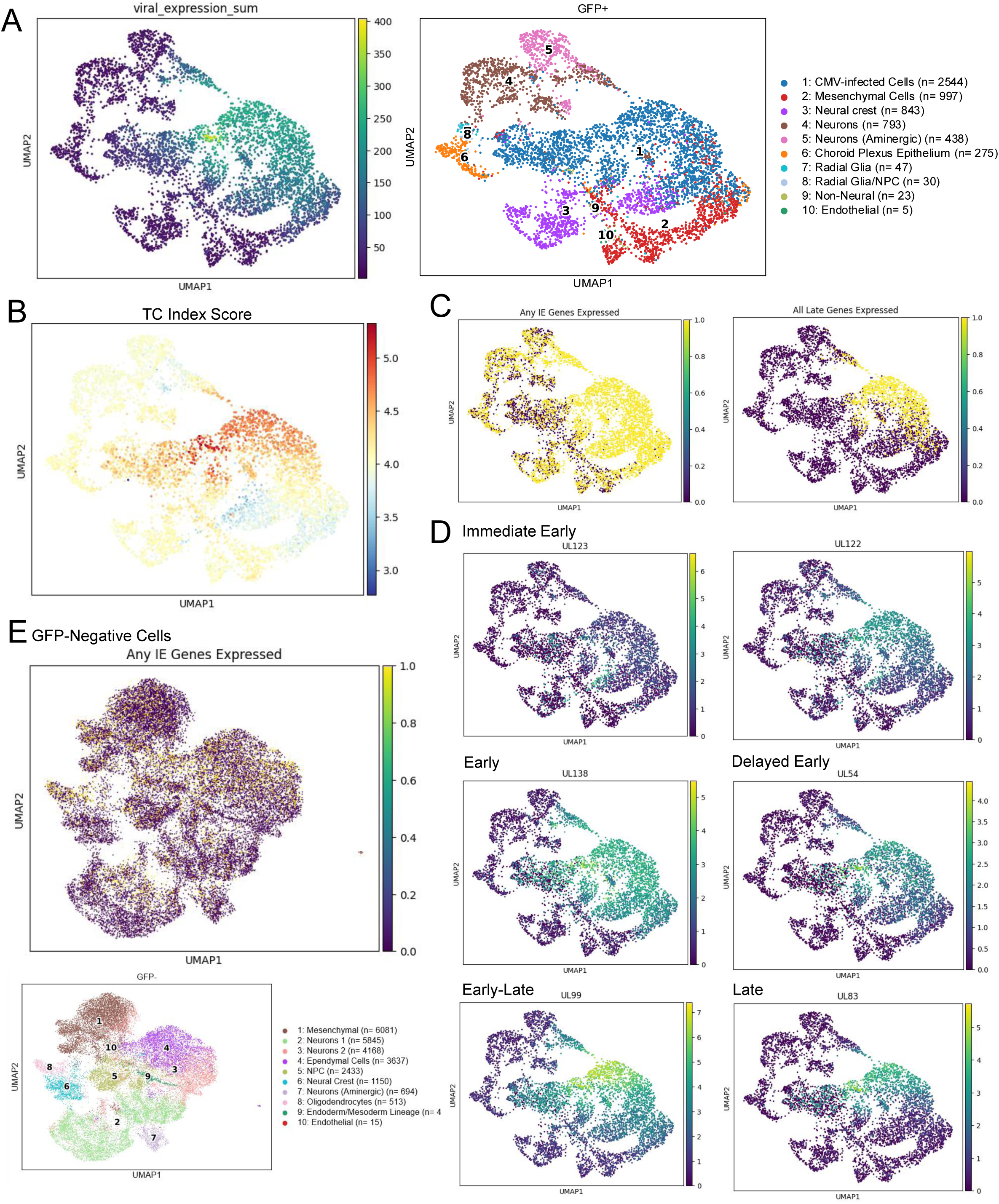
Scoring HCMV-infected cells by temporal progression of viral genes. (A) UMAP plot showing the raw sum of all viral genes across every cell in the GFP-positive subset. UMAP plot of the GFP-positive subset of HCMV-infected neural organoid clustered by cell type using Scanpy (v1.11.5). (B) UMAP density plot showing TC index score across the GFP-positive subset. (C) UMAP plot showing the distribution of cells expressing any HCMV Immediate Early genes (UL122, UL123, IRS1), and UMAP plot showing the distribution of cells expressing all HCMV Late genes (UL32, UL48, UL52, UL94, UL82, UL83) in conjunction. (D) UMAP plots showing HCMV viral gene expression, grouped by the transcriptional infection time point corresponding to each viral gene. (E) UMAP plot showing the distribution of cells expressing any Immediate Early genes in the GFP-negative subset.

To conceptualize the temporal expression of Immediate Early, Early and Late viral genes in each cell, we created a scoring system called TC index score based on the relative balance of module scores for TC1 to TC7 gene groups (**Fig 5B**). Each TC class is composed of many viral genes (**S6A-F Fig**) which can be summarized by modular scores (**S6G Fig**). Low TC index scores correspond to higher relative expression of Immediate Early and Early viral genes (TC1-TC4) compared with a low relative expression of Late viral genes (TC5-TC7). In contrast, higher TC index scores correspond to higher relative expression of Late genes compared with low relative expression of Immediate Early and Early genes. We then scored every cell in the GFP-positive subset along the TC index axis and plotted the scores on a UMAP plot (**Fig 5B**). A large fraction of cells in the CMV-infected Cells (Cluster 1) score high TC index scores (**Figs 5A and 5B; red**). This aggregate cluster expressed high levels of all viral genes (**S6A-F Fig**). With both high UMI counts and TC index score, we postulate that these cells are producing de novo virions. Cells with a low TC index score localized to several UMAP areas (**Figs 5A and 5B; blue**), corresponding to a higher ratio of Immediate Early and Early TC gene expression to late TC gene expression with an overall large number of viral transcripts. This expression pattern is reflected by Mesenchymal Cells (Cluster 2) and Neural crest (Cluster 3) blending into CMV-infected Cells (Cluster 1). These observations suggest a moderate degree of transcriptional similarity between the noted groups while maintaining their host identity markers, unlike the CMV-infected Cells cluster (**Figs 5B and S6A-F Fig**). Since Late gene expression requires viral DNA synthesis, we postulate that a fraction of cells in Clusters 2 and 3 have yet to initiate HCMV DNA replication, evident by differences in GFP expression levels between TC3-only expressing cells and cells expressing TC7 (**S6H Fig**). Finally, several UMAP areas in the GFP-positive population contain cells with intermediate TC index scores (i.e., similar TC1-7 expression levels) along with a range of UMI counts (**Figs 5A** and **5B; yellow**), indicating that these cells are infected yet have failed to initiate the prototypic lytic cycle.

As noted, several clusters in the GFP-positive sample exhibited low total viral gene expression (**Fig 5A**), intermediate TC index scores (**Fig 5B**), and low expressions of individual genes (**S6A-F Fig**). Therefore, we analyzed GFP-positive cells to identify whether or not cells express any IE genes IRS1, UL123, or UL122 (**Fig 5C**). This analysis revealed that 71% of cells are expressing an IE gene. For IE expression levels per cell, we plotted the UMI counts for UL123 and UL122 on the UMAP plot. A range of UL123 expression was observed over the full population with highest levels at the edges of CMV-infected Cell (Cluster 1) and blending with Mesenchymal Cells (Clusters 2) and Neural Crest (Cluster 3). In contrast, we detected the highest expression levels of UL122 in CMV-infected Cell (Cluster 1) (**Fig 5D**) and aligning with a high TC index score (**Fig 5B**). This high score also overlapped with high expression levels (**Fig 5D**) of Early genes UL138 (TC2) and UL54 (TC3), Early-Late gene UL99 (TC4), and Late gene UL82 (TC6) (**S6A-F Fig**). This included a grouping within Cluster 1 expressing all essential Late genes UL32, UL48, UL52, UL94, UL82, and UL83 (**Fig 5C**). As noted earlier, we speculate that these cells are capable of producing *de novo* virus. Finally, we asked whether any IE gene expression could be detected in the GFP-negative cells isolated from the infected organoids (**Fig 5E)**. We observed that 12% of cells that had detectible levels were distributed throughout clusters. These data are consistent with our previous bulk-RNA seq analysis in which IE transcripts were detectable in GFP-negative samples, but with levels significantly lower than GFP-positive samples (19). No Late genes were observed in the GFP-negative cells. Overall, we have demonstrated a distribution of viral gene expressions in cells isolated from HCMV-infected organoids at 14 dpi with varied TC-index scores between clusters. This includes high levels of all viral gene classes and high TC scores in CMV-infected Cells (Cluster 1), but low scores in adjacent Mesenchymal Cells (Cluster 2) and Neural crest (Cluster 3) cells which expressed most TC1 to TC4 genes but had limited expression of Late TC5 to TC7 genes.

### Machine learning classifier elucidates host transcripts that vary with TC index scores

The variation of TC scores in cluster groupings raised the question of whether HCMV infection of tissues with diverse cell types induced altered cells states. This might be the result of delayed or arrested viral replication, which can only be postulated since our analysis did not include a time course of infected tissues. To investigate different possibilities, we binned GFP-positive cells into three quantiles based on their TC progression scores, labeled as low (high TC1 to 4 vs low TC5 to 7), medium (similar TC1 to 7 but varying overall levels), and high (low TC1 to 4 vs high TC5 to 7) bins. We plotted each of these bins on a UMAP (**Fig 6A**). We next sought to use inference-based machine learning to understand what cellular genes are specific to low, medium, or high TC index scores. We trained an XGBoost (eXtreme Gradient Boosting) classifier on the GFP-positive gene expression values to predict the low, medium, or high bin label for every cell. We then generated a confusion matrix (**Fig 6B**) to display discrepancy between true bin label (y-axis) versus the bin label predicted by the model (x-axis). High values along the diagonal signify high accuracy of the classifier model in predicting bin labels. The classifier achieved an overall accuracy of 0.74 (random chance = 0.33) with a macro averaged F1 score of 0.74. The model performed best predicting the high and low classes (F1 = 0.82 and 0.75, respectively) and performed less well in predicting the medium class (F1 = 0.65). These results indicate that the model has learned how to differentiate low, medium, and high TC index labels across cells. To unpack which genes are allowing for accurate classification by the model, we performed SHAP (SHapley Additive exPlanations) analysis and plotted a beeswarm plot for each TC progression bin (**Figs 6C-E**). The beeswarm plot illustrates how various genes’ expression levels (high-red, low-blue) influence the model to predict in a specific direction (right-toward, left-away).

**Fig 6.**
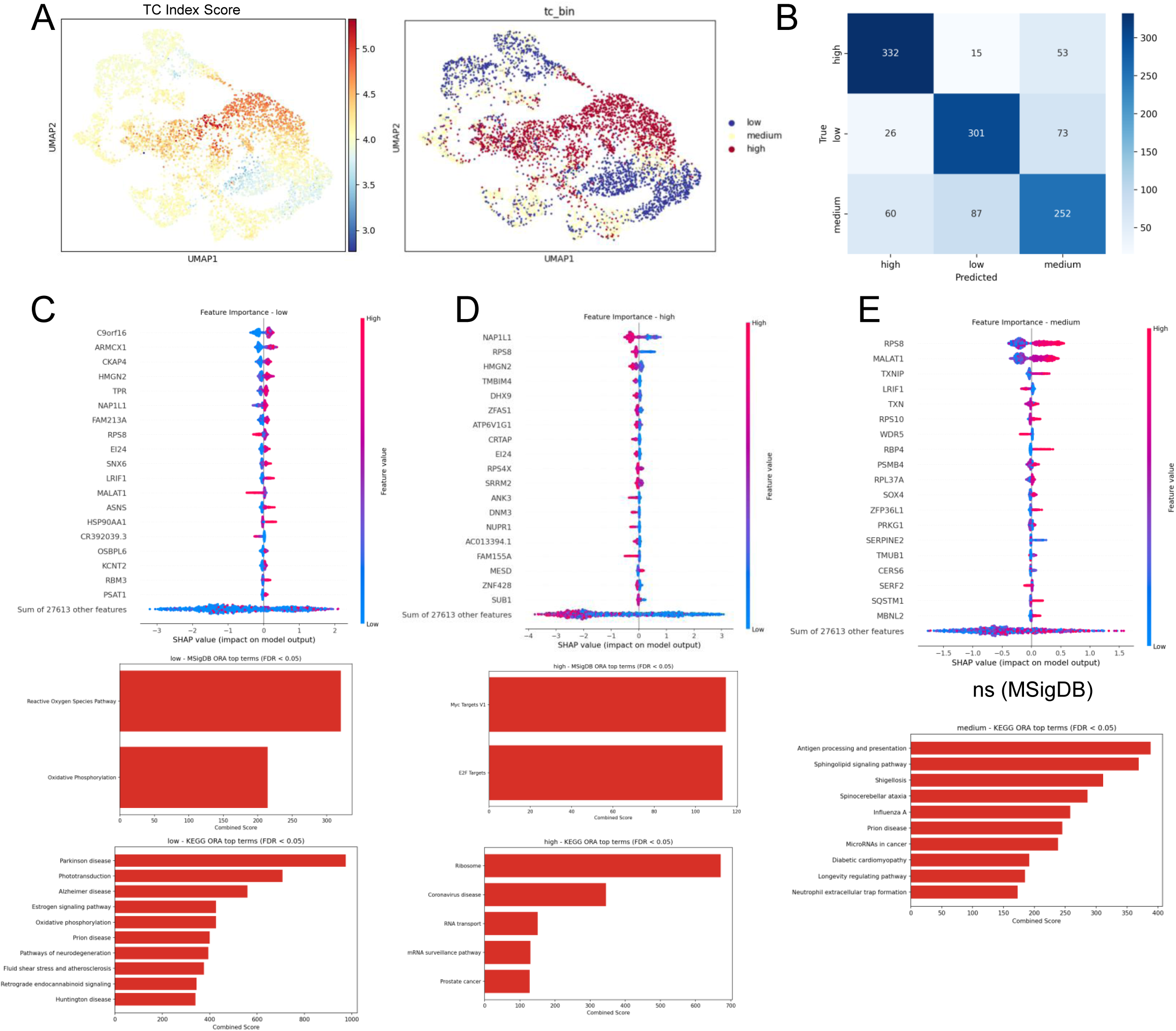
Machine learning inference revealing genes related to HCMV lytic progression. (A) UMAP plot showing a comparison of TC index scores across cells in the GFP-positive subset, and the bin distribution following binning of cells into low, medium, and high quantiles. (B) Confusion matrix for a classifier trained on the expression matrix of the GFP-positive subset to predict low, medium, and high bin labels. (C-E) SHAP beeswarm plots showcasing genes that drive the model to predict a given cell’s label as either toward a given label (to the right) or away from a given label (to the left) for (C) low TC index score, (D) high TC index score, and (E) medium index score. The top 100 positive SHAP genes were used for analysis of MSigDB and KEGG over-representation analysis.

Any cell to the right has information in that cell on a specific gene pushing that cell to that grouping based on TC index (**Figs 6C-E, dots**). Several host gene sets emerge as indicators for TC scores across cell types with accompanying predicted network effects. The low TC progression bin (**Fig 6C; blue**) is defined by cells expressing genes such as C9orf16, ARMCX1, CKAP4, and TPR but not expressing MALAT1, RPS8, and CR392039.3. Notably, cells in this grouping are GFP-positive, expressing a large number of viral genes with high TC1-4 but low TC5-7 Late genes. Using the high impact cellular gene set, we evaluated network effects by performing Over-Representation Analysis. We have included significant terms (FDR<0.05) from both the MSigDB (Molecular Signatures Database) and the KEGG (Kyoto Encyclopedia of Genes and Genomes) databases (**Fig 6C)**. This machine learning approach uncovered terms related to neurodegenerative diseases including Parkinson disease (KEGG), Alzheimer disease (KEGG), Prion Disease (KEGG), Pathways of Neurodegeneration (KEGG), Huntington disease (KEGG), and paralleled by Reactive Oxygen Species Pathway (MSigDB) and Oxidative Phosphorylation (MSigDB). In the high TC index bin (**Fig 6D, red**), which includes the CMV-infected cell Cluster 1, we observe dependence on genes such as NAP1L1, RPS8, and MESD, but not FAM155A, ANK3, and NUPR1. Over-Representation Analysis identifies Myc Targets V1 (MSigDB), E2F Targets (MSigDB), Ribosome (KEGG), and RNA transport (KEGG) which are consistent with known consequences of lytic HCMV replication. Finally, the medium grouping consists of cells with a range of overall viral gene expression but at similar levels of TC1-7 genes (**Fig 6E, yellow**). The wide range of overall levels of viral gene expressions in this grouping makes interpreting the enriched pathways difficult (**Fig 6E**). Overall, while many genes exhibit differential specificity to low, medium, or high TC index scores, it is important to note that the SHAP contribution of any individual gene is significantly outweighed by the contributions of the sum of all the remaining genes not in the plot, indicating that broad-scale network contributions play a more direct role in characterizing HCMV progression from Early to Late TCs.

### Unsupervised hierarchical clustering reveals distinct infection sub-states that overlap with TC index scores

Analyzing cellular states by the TC index is dependent on predetermined viral gene kinetic classes. To evaluate these data without a potential bias, we isolated the HCMV gene expression matrix for every cell and performed agglomerative hierarchical clustering using ward linkage (51). We constructed a dendrogram to visualize clustering to choose optimal cluster number (*k*) (**Fig 7A**). We chose to cut the dendrogram at *k=*7 (silhouette score = 0.13) to get a fine sampling of different viral states present in the scRNA-seq data, and we assigned seven clusters labeled viral sub-states A to G. We plotted a UMAP projection to visualize viral sub-state boundaries (**Fig 7B**) and include the cell type UMAP for comparison. Viral sub-states E to G overlap with the CMV-Infected Cells cluster with sub-state G aligning with the highest level of viral gene expression (**Fig 5A**). The overlap is more evident in a stacked bar plot of cell type composition per each viral sub-state (**Fig 7C**), showing progressively more CMV-infected cells from viral sub-states E to G. Sub-states A to D are adjacent to the CMV-infected cells cluster with sub-state C overlapping with CMV-infected cells and other cell type clusters (**Fig 7B**).

**Fig 7.**
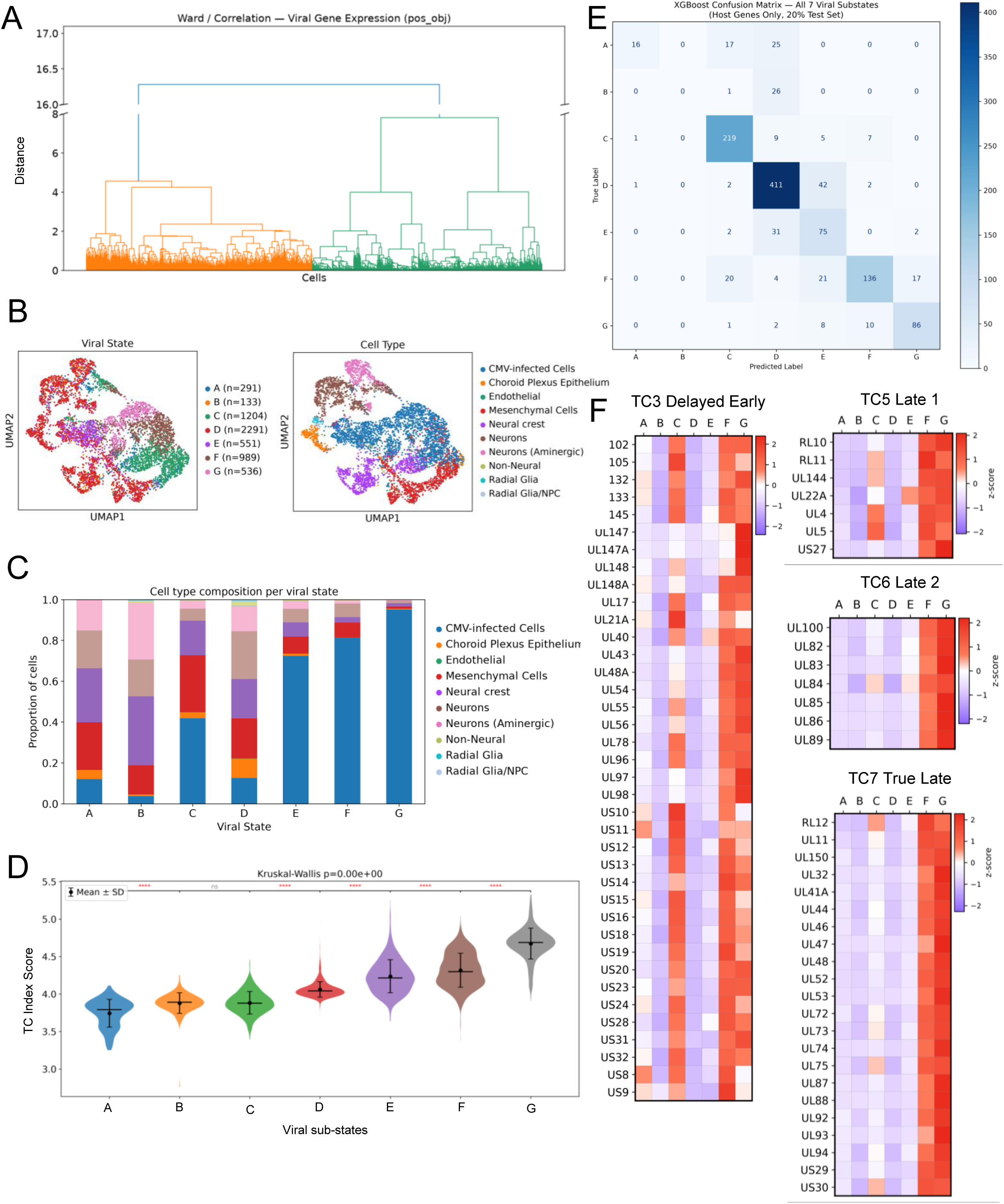
Hierarchical clustering of cells by viral gene expression patterns. (A) Dendrogram derived from unsupervised agglomerative hierarchical clustering using Ward linkage, using only HCMV viral genes and blinded to host genes. (B) UMAP plot showing hierarchical-derived cluster identity (Viral State), compared with UMAP plot showing cell type identity. (C) Stacked bar plot illustrating cell type composition of each Viral State cluster. (D) Violin plot showing distribution of TC index scores grouped by Viral State cluster. Inter-cluster statistical significance was calculated using the Mann-Whitney U test. (E) Confusion matrix for a model trained on the host gene expression matrix to predict cluster labels. (F) Heatmaps of each viral state’s average expression of HCMV genes, grouped by temporal class Early TC3 genes and Late TC5-7 genes.

To evaluate whether viral sub-states A to G correlate with the TC index scores, we created a violin plot of index scores for the cells comprising each viral sub-state (**Fig 7D**). The adjacent viral sub-states show a consistently significant increase in TC index score by pairwise Mann-Whitney U (Kruskal-Wallis H test: H=3828.93, *p*∼0). Notably, viral sub-state C, despite having a higher overall expression of viral genes than sub-state D, has a lower TC index score (**Fig 7C**). This is a result of sub-state D expressing proportionally more Late viral genes than it expresses Early viral genes, albeit at an overall low level, shifting the balance toward a medium TC index score. This is a reflection of Softmax scoring within the TC index calculation, which reduces absolute viral expression levels to expression levels relative to the other viral genes that a given cell is expressing.

Considering the low silhouette score of *k*, we sought to confirm that the viral sub-states assigned at *k=7* were useful for analyses and that it captured unique viral sub-states. To accomplish this validation, we trained an eXtreme Gradient Boosting (XGBoost) classifier to label cells by their viral sub-state using only host genes as features (**Fig 7E**). Across all viral sub-states and all cells in the GFP-positive subset, the model achieved a ROC-AUC score of 0.936, predicting viral sub-states C to G with a high degree of accuracy. Together with the silhouette scores, these data suggest a more-continuous gradient across viral sub-states in the dimension of increasing TC index, rather than perfectly quantized states approximated by the clustering. Viral sub-states A and B were poorly differentiated by the classifier model, likely due to high preservation of cell type identity in lower infected clusters, obscuring the shared transcriptional program activation present across cell types in highly infected clusters. Additionally, small cell numbers likely contributed to class imbalance during training, limiting classification fidelity toward those states.

Finally, we aligned each viral gene class to the uncovered viral sub-states for A to G. We plotted a heatmap of average viral gene expression against each viral sub-state (**Figs 7F and S7**). The sub-states F and G exhibit high expression across all viral genes including Late genes. Sub-states A and B exhibit much lower expression. Sub-states A and B exhibit expression of UL123 and UL122, with sub-state A expressing higher overall levels of TC1, TC2 and TC3 gene classes (**S7A-C Fig**). Viral sub-state D is characterized by very low expression of all viral genes, which we interpret to represent cells that have not meaningfully entered the HCMV lytic progression despite expressing GFP. Viral sub-state C is an outlier exhibiting a scattered distribution of viral gene expression, with a variety of different transcripts being expressed with moderate bias toward expression of TC3 viral genes. This observation further points toward viral sub-state C representing a distinct state that overlaps clusters of HCMV-infected cells (Cluster 1) (i.e., lost identity markers) with Mesenchymal Cells (Cluster 2) and Neural Crest (Cluster 3) cells.

### Analysis of host genes driving different viral sub-states reveals network dysregulation

After analyzing the data from this different perspective, we re-examined host cellular processes altered between viral sub-states A to G. We performed differential expression (DE) analysis to compare transcriptional makeup of each viral sub-state with all other cells in the GFP-positive population, then plotted the top 10 marker genes for each sub-state on a dot plot (**Fig 8A**). For example, sub-state C consists of a high percentage of cells expressing genes SRM, MANF, and TOMM40 compared to all other sub-states, most notably sub-state E. To analyze pathway-level patterns of expression within each viral sub-state, we performed preranked Gene Set Enrichment Analysis (GSEA) on differentially expressed host genes for each viral substate, then thresholded top terms by a False Discovery Rate (FDR) threshold of <0.1 and plotted positive normalized enrichment score (NES) terms that were differentially enriched in each viral state (**Fig 8B**). Sub-states B and D, both of which have low levels of viral gene expressions, are significantly enriched in genes involved in several processes including interferon responses, KRAS signaling, and the epithelial mesenchymal transition (EMT). In contrast, sub-states E and F are enriched in genes associated with EMT and reactive oxygen species pathways, respectively. Of particular interest again is viral sub-state C, which exhibits high TC1 to 4 but low TC5 to 7 gene expression. We observe enriched genes associated with E2F and Myc gene targets, mTORC1 and the unfolded protein response, glycolysis and adipogenesis, and G2-M checkpoint. All of these processes are known targets of HCMV protein function, occurring early during viral infection when studied in monolayer culture systems.

**Fig 8.**
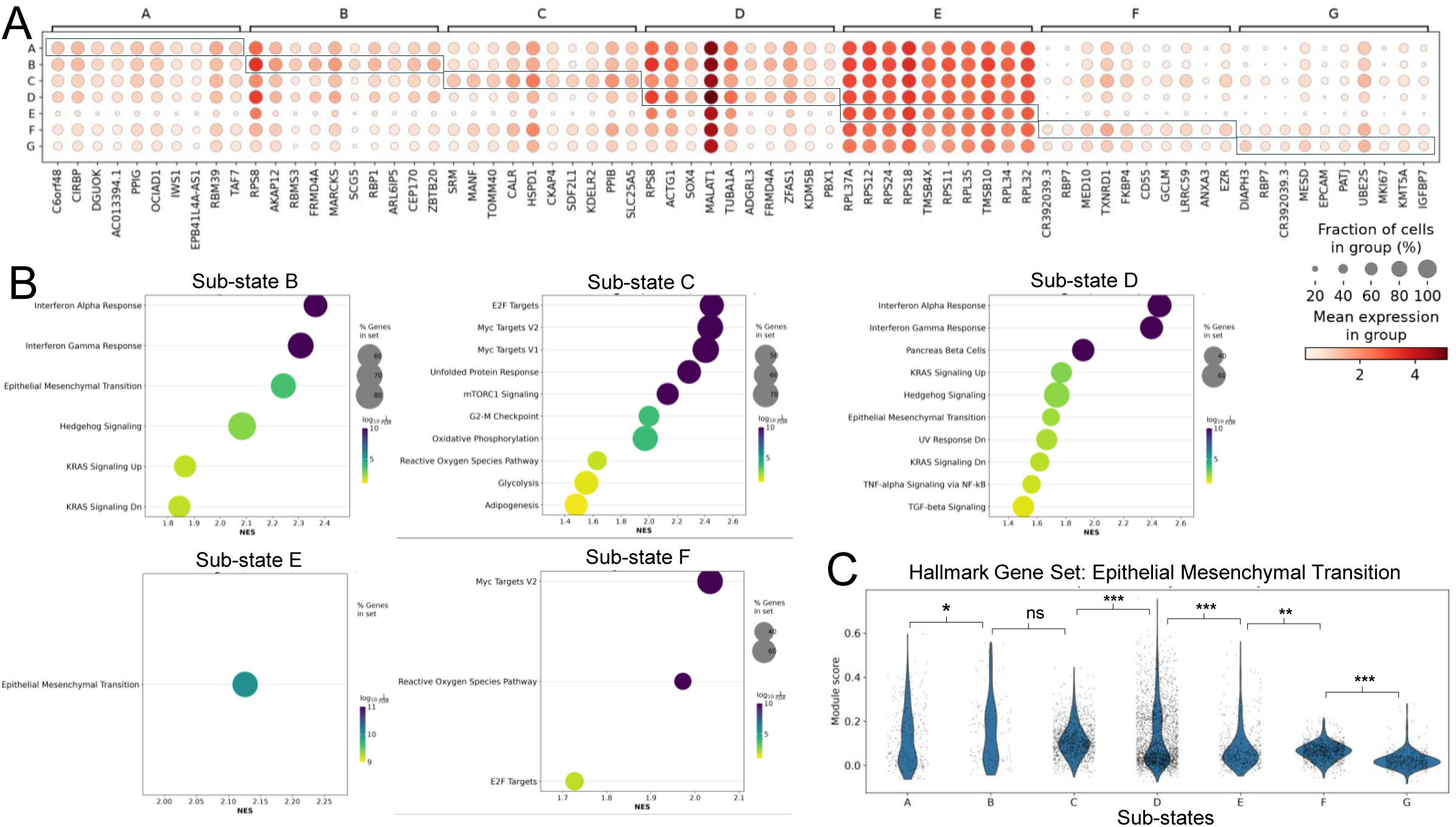
Differential host gene responses to different viral states elucidated from hierarchical clustering. (A) Dot plot of top 10 differentially expressed host genes in each viral sub-state cluster. (B) Plot of significant terms (FDR<0.1) from pre-ranked Gene Set Enrichment Analysis (GSEA) of differentially expressed genes in each viral sub-state cluster. GSEA was constructed by querying the MSigDB Hallmark 2020 database. (C) Violin plot showing module scores of Epithelial to Mesenchymal Transition (EMT) genes from the MSigDB Human gene set HALLMARK_EPITHELIAL_MESENCHYMAL_TRANSITION, grouped by viral sub-state cluster.

One Hallmark Gene set enriched in several sub-states is the Epithelial Mesenchymal Transition (EMT). Several studies using varying cell types have implicated EMT or the reverse transition during HCMV infection. In the GFP-positive cell population in infected organoids, EMT was significantly overrepresented (NES>0) in viral sub-states B, D, and E, but was underrepresented (NES<0) in viral sub-states F and G. We analyzed each sub-state by generating a modular score per cell for EMT hallmark genes (**Fig 8C**). We observe varying modular scores for the EMT gene set that are significantly different between sub-states with the score elevated in viral sub-state C (Mesenchymal (Cluster 2), Neural Crest (Cluster 3), low TC index score with high levels of viral gene expression) and decreasing in sub-states as they progress to sub-state G (CMV-infected Cell Cluster 1, high TC index score, high levels of viral gene expression).

### Analysis of EpCAM Expression *in vitro* reflects downregulation observed in scRNA-seq

One notable gene associated with EMT is EpCAM (Epithelial Cell Adhesion Molecule, CD326). We plotted EpCAM expression by viral sub-state on a violin plot and observed a significant upregulation of EpCAM, which appears as one of the top differentially expressed genes (DEGs) in viral sub-state G (**Figs 9A and 9B**) and notable expression in viral sub-states C, D and F. We plotted EpCAM expression density on a UMAP plot to visualize expression and observed a higher concentration of EpCAM in the clustering of cells likely to be producing *de novo* virus (**Fig 9B**). We have defined these cells as CMV-infected Cell Cluster 1, high TC index score, and viral sub-state G. EpCAM expression is not unique to the GFP-positive cells and is elevated in several clusters in Mock-treated (Clusters 11-14) and GFP-negative (Cluster 9) cell populations (**S9 Fig**). Our lineage fate analysis indicates that GFP-negative Cluster 9 originates from Mock Cluster 13 (**Fig 4A**), and the CMV-infected cell Cluster 1 partially originates from Mock Clusters 11-14 (**Fig 4B**). Therefore, it is not clear whether infection is inducing expression or whether HCMV infects cells with elevated EpCAM. Previous studies demonstrated infection-induced EpCAM expression in epithelial and fibroblasts monolayer cultures at late times of infection (66,67).

**Fig 9.**
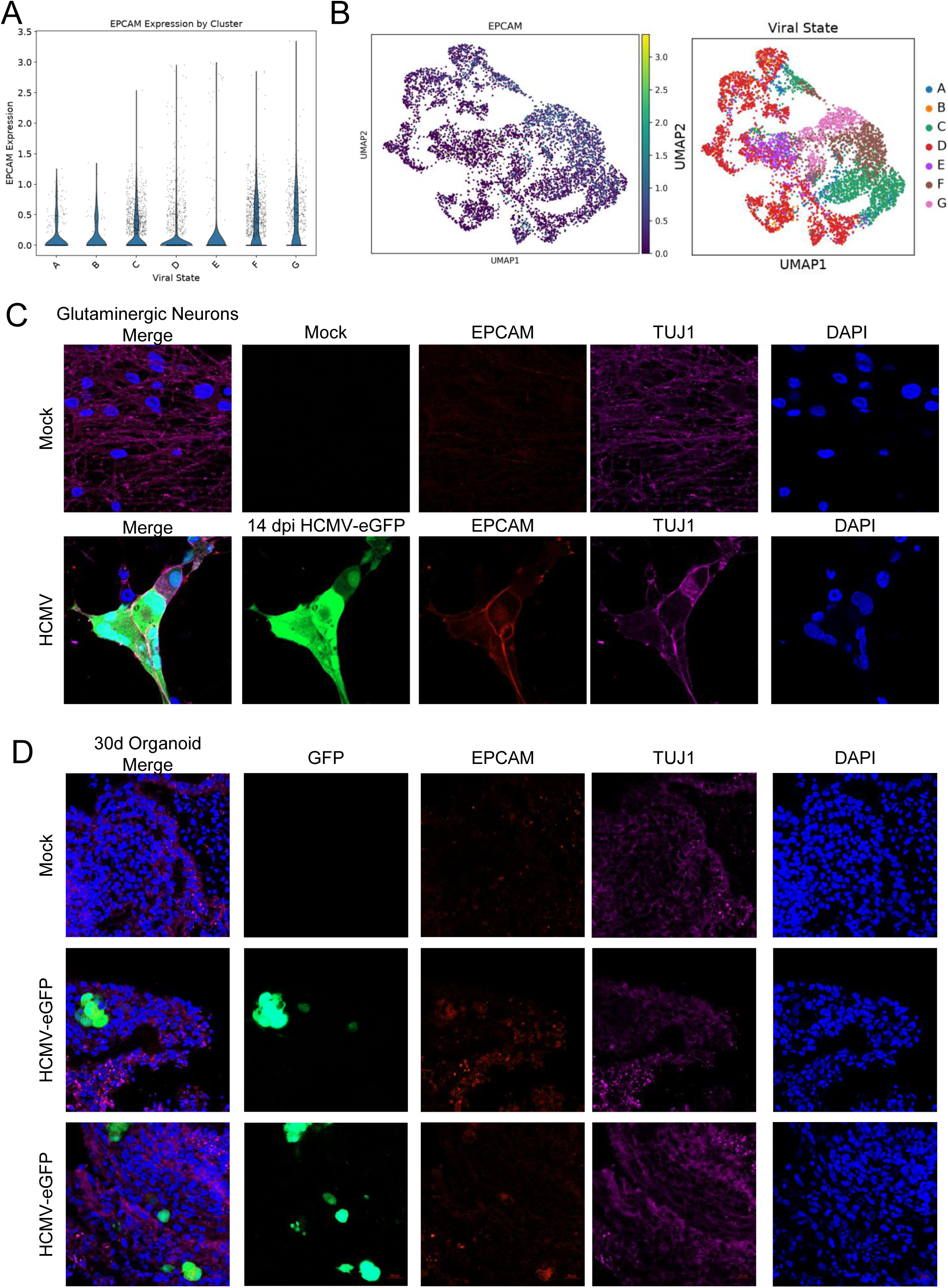
EpCAM expression in HCMV infected glutamatergic neurons and cortical organoids. **(A)** EpCAM expression in different viral sub-states shows of the GFP-positive populations. (B) UMAP plot of EpCAM expression and viral sub-states in the GFP-positive subset of the HCMV-infected cortical organoid. (C) Immunofluorescence images of mock and HCMV-infected (eGFP green) human glutamatergic neurons stained for EpCAM (red), neuronal marker TUJ1 (purple), and DAPI (blue). (D) Immunofluorescence images of mock and HCMV-infected (eGFP) human cortical organoids with staining described in (C).

To experimentally validate the information obtained by our analyses, we sought to examine EpCAM expression in terminally differentiated neurons and cortical organoids. We infected 45-day old iPSC-derived cortical neurons and 30-day old iPSC-derived cortical organoids with HCMV TB40 eGFP. At 14 dpi, we performed immunofluorescence analysis staining for EpCAM and the neuronal marker TUJ1 (betaIII-tubulin). Mock-treated neurons exhibit robust staining for TUJ1 with little to no detectable EpCAM expression (**Fig 9C**). In contrast, we observed strong EpCAM intracellular and surface staining in HCMV infected cells (**Fig 9C**). These data demonstrate that infection is inducing EpCAM expression in terminally differentiated neurons. In mock-treated neural organoid tissues, low levels of EpCAM are detectable in tissue sections (**Fig 9D**) which increases upon HCMV infection at 14 dpi (**Fig 9D**). The increase is visible in GFP-positive and GFP-negative cells with GFP-positive cells exhibiting a distinct EpCAM staining pattern. These data together demonstrate that HCMV infection is inducing EpCAM expression in neurons and neural organoids.

## DISCUSSION

Congenital HCMV infection is the leading viral cause of birth defects and can result in significant damage to the CNS during development. In the present study, we conducted scRNA-seq analysis of HCMV-infected cortical organoids to determine how infection might impact the developing brain. In 44 day old mock-treated cortical organoids, we identified 16 cell type clusters based on gene expression signatures, with glutamatergic neurons, forebrain NPCs, and mesenchymal cells comprising the largest populations (**Figs 1 and S1**). We used spatial transcriptomics to elucidate changes in cell type composition and distribution in the cortical organoids (**Fig 3**); however, no significant changes were detected following infection, likely attributed to the limited tissue size and probe set. After exposing tissues to HCMV encoding GFP, at 14 dpi we separated cells into GFP-positive and -negative populations by FACS for scRNA-seq and analyses. In the GFP-negative population, low levels of HCMV Immediate Early genes but no Late genes were detected in a fraction of the cells, which is consistent with our previous studies (**Fig 5E**) (19). The most abundant clusters in the GFP-negative population included mesenchymal, neurons, and ependymal cells (**Figs 2B and S2**). Comparative analysis of GFP-negative groupings to mock suggests that cells largely maintain their identity between mock and GFP-negative (**Figs 4 and S4 Fig**).

In the GFP-positive populations, cell types included mesenchymal cells, neural crest cells, and a cell type cluster we labeled “CMV-infected cells” due to an abundance of viral genes and reduced host gene signatures (**Figs 2 and S3 Fig**). The gene signatures persisting in cells of the CMV-infected cell cluster potentially originate from glutamatergic neurons, hindbrain NPCs, and radial glia/NPC (**Figs 3 and S4 Fig**). Interestingly, we did not identify glutamatergic neuron gene signatures in infected organoids, which is consistent with our previous studies showing that this cell type is fully permissive to HCMV lytic replication (12). In contrast, identities of aminergic neurons were maintained following infection with HCMV. In summary, these studies suggest several cell type identities are conserved following infection (e.g., Aminergic Neurons, Neural Crest) whereas others are altered or lost (e.g., Glutamatergic/GABAergic Neurons, Radial Glia).

Cortical organoids offer a unique model system to study the impact of HCMV infection on neurodevelopment (15,16,18–22). In transitioning from monolayer to three-dimensional and multicellular cultures, we have had to address several challenges related to experimental design and asynchronous infections. To study infection in organoids, the field currently uses a variety of methods including variations in inoculum (e.g., viral genotype, concentration, exposure time, etc.) and tissue features (e.g., iPSC source, developmental time, etc.), which make direct comparisons challenging. Despite these differences, previous studies consistently follow the Lancaster and Knoblich protocol (38). For our studies, we used BAC-derived virus, originating from the TB40/E virus (39), produced in epithelial cells, and titered using a limiting dilution TCID_50_ assay. We infected each organoid for 2 hours based on weight at 500 IU/µg at 30 days with tissues generated using PBMC-derived iPSCs and a commercial organoid kit based on Lancaster and Knoblich (38), then analyzed at 14 dpi (**Figs 1A and 2**). Studies by Brown et al. (22) used neonatal brain tissue-derived iPSCs, infected using HCMV clinical virus TR generated in fibroblasts at 5 PFU/cell, and then made organoids out to 52 days. In contrast, Sun et al. (20) used human fibroblasts (old and young)-derived iPSCs, infected using BAC-TB40 (in epithelial) and BAC-Towne (in fibroblasts) at 45 day with 5×10^5^ PFU in 1 ml for 24 hours and analyzed at several times to 20 dpi. Finally, studies by Egilmezer *et al.* (15) used cord blood-derived iPSC and infected at 55 days using Merlin strain at 1×10^7^ PFU for 24 hours and analyzed at several times to 14 dpi. Despite experimental differences, all these studies demonstrate that HCMV infects iPSC-derived neural organoids, disrupts tissue organization including rosette formation, and reduces neurodevelopmental gene expressions by 14 dpi. In addition, we and others have demonstrated that anti-HCMV neutralizing antibodies can limit tissue damage (16,20).

A second challenge in studying infection of organoids is in how to define progression of HCMV replication in a three-dimensional tissue that is inherently asynchronous as the virus spreads to different cell types. The field has defined a linear progression of Immediate Early, Early, Early-Late, and Late gene expressions following a synchronized infection of monolayer cultures, which has been refined using RNA-seq into Temporal Classes TC1 to 7 (35,45). After 14 dpi, defining the state of infection is challenging. Therefore, we introduced a TC indexing approach to define replication status in scRNA-seq data. Each cell is scored based on the relative balance of module scores for the TC1 to TC7 gene groups (**Fig 5B**). In other words, the index score corresponds to the transcriptional progression from expression of Immediate Early TC genes (TC1) to True Late TC genes (TC7). We show that cells with the highest TC score (**Fig. 5B)** overlap with cells in the CMV-infected cell cluster (**Fig 5A**), highest UMI values (**Fig 5A**), expression of the true late gene UL83 (**Fig 5D**), and viral sub-state G (**Fig 7B**). We hypothesize that these cells are actively producing new virus in the tissues. In contrast, the low index scored cells have high levels of Immediate Early and Early genes, for example UL138 (**Fig 5D**), but little Late gene expression, such as UL83 (**Fig. 5D**). We postulate that cells in this grouping have yet to initiate viral DNA synthesis. One limitation we encountered was the presence of cells with intermediate index scores (i.e., similar levels of TC expressions) but varying overall UMI levels, requiring future method refinement. Overall, the use of TC indexing has allowed us to study varying HCMV-induced cell states in an asynchronous manner.

To further analyze asynchronous cell states, we introduced machine learning techniques including classifiers (**Fig 6**) and unsupervised clustering (**Fig 7**) to analyze nonlinear relationships and higher-order interactions. Using unsupervised clustering to characterize clustered snapshots of viral infection (**Fig 7**), we observed pathway-level changes across cell types in response to variable expression of viral genes (**Fig 8**). Such usage of unsupervised machine learning methods is useful in detecting lower-dimensional gene programs important to the fundamental biology of a system. Interpreting fewer dimensions, each comprised of many component genes, allows for the preservation of complexity in the data itself while providing the interpreter with greater simplicity. We also used supervised methods such as machine learning classifiers to elucidate unifying factors across cell types that underlie HCMV progression (**Fig 6**). This revealed the importance of host systems of inflammation, oxidative stress, and even more subtle changes including ribosomal subunit usage across infection course.

We observed that HCMV infection of neural organoids induces altered cell states and found evidence suggesting an epithelial to mesenchymal-like (EMT) transition. EMT generates cells with mesenchymal markers and increased motility, participating in processes such as embryogenesis, wound healing, and cancer progression (52). Interestingly, cells infected by HCMV have been shown to have increased motility (53–56). Using unsupervised clustering of cells based on viral gene expression patterns in the GFP-positive population, we observed enriched gene sets defining EMT in several viral sub-states (**Fig 8**). The highest modular scores for EMT genes are found in sub-state C and D, and the score is decreased in sub-state G which contains the CMV-infected Cell cluster. These sub-states also exhibited elevated levels of EpCAM (**Fig 9A**), which we validated at the protein level using infected neurons and organoid tissues (**Figs 9C and 9D**). EpCAM is highly expressed on epithelial tumors and demonstrated to be an effective target of CAR-T cells (57,58). This further raises the possibilities of CAR-T or vCTL-based strategies against refractory HCMV disease (59–61). The role of HCMV infection in cancer is controversial. However, several laboratories have implicated EMT, or its reverse MET, following infection. HCMV infection of renal tubular epithelial cells induces EMT involving TGF-β (62) which was also observed following infection of iPSC-derived cholangiocyte-like cell organoids (63). Induction of TGF-β in cells is mediated by HCMV UL123 (IE1 protein) and UL122 (IE2) (62,64,65). We observed significant enrichment of TGF-β hallmark genes in viral sub-state D (**Fig 8D**), which exhibits the highest modular score for EMT accompanied by low overall levels of viral gene expressions (**S7 Fig**). In contrast, studies using retinal pigment epithelial cells demonstrate HCMV-induced MET with increased EpCAM expression at late times of infection (66,67). Regardless of the direction of transition, it is clear that infection is inducing unique cellular sub-states prior to the HCMV-infected Cells cluster that lacks sufficient information to define their developmental states. Importantly, our studies use organoid cultures that involve a neuroepithelial expansion phase. Recent studies using fibroblasts have demonstrated that HCMV infection also induces neuronal gene signatures including enolase and macroH2A1 (40,68,69). Together, these observations raise questions as to whether HCMV proteins reprogram cells to neuroepithelial-like states to support fulminant infection, and whether this conducive state explains the detection of HCMV in glioblastoma tumor biopsies.

Our hierarchical clustering approach also identified a sub-state of particular interest, viral sub-state C, that expresses Early genes but few Late genes (low TC index score) (**Figs 5B, 7B and 7C**). There is a scattered distribution of specific viral genes, with moderate bias toward expression of viral genes defined as Delayed Early or TC3 genes (**Figs 7F and S7 Fig**). The low levels of Late gene (TC6, 7) and GFP expression (**S6 Fig**) suggest that HCMV genome replication has not occurred in the population. Cells in this classification overlap with clusters in the CMV-infected cells (Cluster 1) (that have lost identity markers), Mesenchymal Cells (Cluster 2), and Neural Crest (Cluster 3) cells (**Fig 7B**), including cells expressing high levels of EpCAM (**Figs 9A and 9B**). Differentially expressed cellular genes in viral sub-state C implicate increases in E2F and Myc target genes, the unfolded protein response (UPR), mTORC1 signaling, G2-M checkpoint, oxidative phosphorylation, glycolysis and adipogenesis (**Fig 8**). HCMV UL122 (IE2), UL97, and UL82 have been demonstrated to inactivate RB and stimulate E2F gene expression (70–77). Intermediate levels of these genes are observed in sub-state C (**S7 Fig**). We also observed increased signatures associated with mTORC1 regulation, which has been previously attributed to expression of HCMV UL38 and its disruption of cellular tubular sclerosis complex 2 (TSC2) (78–81). Expression of UL38 is at its highest level in cells within sub-state C of infected tissues (**S7 Fig**). Infection, including expression of UL38, stimulates glycolysis and lipogenesis (82–88). Finally, the UPR is induced during HCMV infection, stimulated by UL148 (89–91), which is also expressed at high levels in sub-state C (**S7 Fig**). Overall, our strategy in grouping cells uncovered a unique viral sub-state C, exhibiting altered cellular processes with defined mechanisms by the expressed viral genes. It is not clear whether these cells can progress to support HCMV genome replication as determined by GFP levels and late gene expression. Although a lofty goal, we are interested in defining which viral genes might be sufficient to construct this unique viral-sub-state in cortical tissues. Failure of infection to progress to viral genome synthesis and lysis may result in a cellular sub-state with hallmarks of a cancer cell.

In summary, our studies using scRNA-seq of HCMV infection of multicellular three-dimensional cortical organoids revealed unique features of infection, including varying viral sub-states and tropisms, and introduced new approaches to analyze complex data sets consisting of cells in multiple developmental states and various stages of HCMV infection.

## MATERIALS AND METHODS

### iPSC cortical organoid differentiation and infection

Cerebral organoid cultures were differentiated from a healthy iPSC line HB53 according to manufacturer instructions (STEMCELL Technologies (#08570)) Briefly, iPSCs were seeded at 9×10^3^ cells per well onto 96-well ultra-low attachment plates for embryoid body (EB) formation and grown in EB formation media (STEMCELL Technologies). On day 5, induction of neural epithelium was initiated by moving the EBs into an ultra-low attachment 24-well plate where they were then fed with induction media (STEMCELL Technologies). On day 7, neural tissues were embedded in Matrigel droplets and moved to ultra-low attachment 6-well plates and fed expansion media (STEMCELL Technologies). On day 10, the plate of developing organoids was transferred to a rocker to circulate nutrients and prevent organoids from attaching to the plate. From this point on, the organoids were fed maturation media (STEMCELL Technologies) every 3 days. On day 30, organoids were weighed and infected with 500 infectious units/µg BAC-derived HCMV strain TB40 encoding GFP for 2 h then virus was removed, and fresh media added. After infection, media was then changed every 3 to 4 days until 14 days post infection (dpi). Viral stocks were generated on ARPE-19 cells, concentrated by pelleting through a sorbitol (20% sorbitol, 50 mM Tris-HCl, pH 7.2, 1 mM MgCl_2_) cushion at 20,000 x g for 1 h in a Sorvall WX-90 ultracentrifuge and SureSpin 630 rotor (Thermo Fisher Scientific), and titered using a using a limiting dilution assay (TCID_50_). GFP-positive wells were determined at 2 weeks post infection and resulting titers were referred to as infectious units per milliliter (IU/mL).

### Cortical neuron differentiation

Healthy iPSC line HB53 was differentiated into forebrain glutamatergic neurons as we have done previously. Briefly, iPSCs were patterned toward neural progenitor cells using dual SMAD inhibition (STEMdiff SMADi Neural Induction Kit, STEMCELL Technologies) followed by patterning toward forebrain neurons (STEMdiff Forebrain Neuron Differentiation Kit, STEMCELL Technologies). Neurons were maintained and matured in STEMdiff Forebrain Neuron Maturation Medium (STEMCELL Technologies) until use. After 45 days of culture, neurons were infected with HCMV strain TB40/E encoding GFP at an MOI of 3 for 2 h. Virus was removed, and fresh media added. Cells were fixed and analyzed at 14dpi.

### Organoid sample preparation and scRNA-seq

Individual iPSC-derived cortical organoids were dissociated into a single-cell suspension using the Worthington Papain Dissociation System kit (Worthington Biochemical). Dissociated cells were resuspended in ice-cold PBS containing 0.04% BSA (Sigma) and FACs sorted based on GFP expression into either a GFP (High) or GFP (Low) versus Mock infected as done previously (19). After sorting, cell populations were then resuspended in ice-cold PBS containing 0.04% BSA (Sigma) at a concentration of 1,000 cells/μl, and approximately 17,400 cells per channel (to give estimated recovery of 10,000 cells per channel) were loaded onto a Chromium Single Cell 3′ Chip (10x Genomics, PN-120236) and processed through the Chromium controller to generate single-cell gel beads in emulsion (GEMs). scRNA-seq libraries were prepared with the Chromium Single Cell 3′ Library & Gel Bead Kit v.2 (10x Genomics, PN-120237). Libraries from different samples were pooled based on molar concentrations and sequenced on a NextSeq 500 instrument (Illumina) with 26 bases for read 1, 57 bases for read 2 and 8 bases for index 1. After the first round of sequencing, libraries were re-pooled on the basis of the actual number of cells in each and re-sequenced to give an equal number of reads per cell in each sample and to reach a sequencing saturation of at least 50% (in most cases >70%). After generation of the pooled libraries, libraries were sent for sequencing by Novogene.

### Analyses of the scRNA-seq data

#### Alignment, and Filtering and Normalization

Following library preparation and sequencing, FASTQ files from the Mock, EPIpos, and EPIneg samples were provided for data analysis. A custom joint human–HCMV reference was constructed by combining the human GRCh38 reference FASTA and GTF files (GTF genome build GRCh38.p12) with the HCMV genome.fa and genome.gtf files obtained from the HCMV reference-genomes repository. The viral reference corresponds to human cytomegalovirus strain TB40/E, clone TB40-BAC4 (GenBank accession EF999921.1), and includes the GFP sequence and annotation. The combined human and viral FASTA and GTF files were indexed using software Cell Ranger (v 9.0.0) with command cellranger mkref. FASTQ files from each sample were then aligned independently against the customized human–HCMV reference, using command cellranger count. The filtered gene–barcode matrices were imported into R package Seurat (v5.4) as three independent objects. For quality control, features detected in fewer than three cells were excluded. Cells were excluded if they expressed fewer than 200 or more than 10,000 genes, or had mitochondrial transcript percentages greater than 20%.

Each object was normalized independently using Seurat’s LogNormalize method with a default scale factor of 10,000. Highly variable features were selected using the variance-stabilizing transformation (vst) method. For the EPIpos and EPIneg objects, an HCMV expression module score was calculated (using R function AddModuleScore) from the viral genes in the custom annotation, and mitochondrial transcript percentage and the HCMV module score were regressed out during data scaling step. Principal component analysis was performed independently for each object, and top 30 components were used as input for UMAP algorithm to generate two-dimensional embeddings.

#### Cell Type Cluster Annotation

Specific cell type markers from He *et al*. 2024 and Velasco *et al.* 2019 were plotted by violin plot across all cell type clusters to check for unique upregulation within a given cluster compared to other clusters (40,41). Clusters with clear expression of a given cell type’s marker genes were annotated by that cell type. Remaining clusters of ambiguous identity were subsequently annotated by inputting the top 20 marker genes derived from Seurat’s FindAllMarkers() through AnnDictionary (42) to programmatically query Claude Haiku 4.5. Responses were then used to annotate the remaining cell type clusters.

#### Optimal Transport Analysis

Optimal Transport analysis was conducted using Multi-Omics Single-Cell Optimal Transport (MOSCOT) (v0.5.0) (48). To perform this analysis in Python (v3.11.9), we exported the Mock and GFP positive Seurat objects’ normalized expression matrices, cell metadata including cell type annotations, gene metadata, and raw counts as h5ad files. H5ad files were then read into AnnData (v0.12.16) objects, which we used for all subsequent analyses in Python. We next concatenated these objects to create a combined Mock and GFP positive AnnData object. We subsequently removed HCMV viral genes from the concatenated AnnData object, and computed top 3000 highly variable genes, performed a PCA dimensionality reduction to 30 components, computed k nearest neighbors, and plotted using 2-dimensional UMAP all using Scanpy (v1.11.5).

MOSCOT analysis was conducted solving the TemporalProblem (49) setting Mock time_key to 0 and GFP positive time_key to 1. This was done with the objective of modeling HCMV infection as a temporal perturbation to determine probabilistic transcriptional mapping of Mock cell type clusters onto GFP positive cell type clusters following infection. Solution of the temporal problem was called using parameters tau_a=0.95 and tau_b=0.90. These parameters were set to reflect unbalanced optimal transport, as we sought to account for the loss of some transcriptional identity following HCMV infection, as evidenced by the presence of a large UMAP cluster of CMV-infected cells whose markers were comprised predominantly of HCMV viral gene transcripts. Following solution of the TemporalProblem, we constructed a Transition Probability Matrix heatmap to illustrate the probability mass of cells within Mock clusters transitioning to cells within GFP positive clusters. We additionally constructed a Sankey plot to illustrate flow of probability mass between Mock and GFP positive cell type clusters. We thresholded the Sankey plot to transition probabilities greater than 15% to enhance visualization of network patterns in the plot.

### Sample Preparation of Organoids for Spatial Transcriptomics

Individual iPSC-derived organoids were prepared for cryosectioning following an adaptation of the Visium CytAssist Spatial Gene Expression for Fixed Frozen – Tissue Preparation Guide (CG000663 Rev A) as suggested by the manufacturer. In brief, organoids were fixed in 4% paraformaldehyde (PFA) overnight, cryoprotected in 30% sucrose overnight, embedded in optimal cutting temperature (OCT), and then flash frozen utilizing an isopentane bath (Millipore Sigma, M32631). Fixed-frozen OCT-embedded tissue blocks contained 5 organoids (3 HCMV, 2 mock) in a row. Organoid sectioning was performed following the 10X Genomics Protocol CG000579 Rev F. Prior to sectioning, Xenium slides were equilibrated to the CryoChamber for 30 minutes. Tissue was sectioned at 10μm thickness and placed on a Xenium Slide (10X Genomics) according to manufacturer protocol. Three serial sections with approximately 30uM between each section were placed on each slide, with two slides total consisting of 10 different organoids (6 HCMV, 4 mock). Slides were stored in at −80°C until sample preparation.

For rehydration and decrosslinking, the 10X Genomics Protocol CG000662 Rev B using Xenium Sample Prep Reagents (10X Genomics, 000460) was followed with minor adaptations according to manufacturer’s suggestion. For the remainder of sample preparation, slides were processed through hybridization, ligation, amplification, and segmentation protocols (10X Genomics Protocol CG000749 Rev B). In brief, the pre-designed Xenium Human Brain Gene Expression Panel (10X Genomics, 1000599) containing 266 gene probes were added to the tissue, hybridized overnight, and then ligated at 37C for 2 hours. Following ligation, probes were amplified for 2 hours at 30C. For cell segmentation using the Xenium Cell Segmentation Add-on Kit (10X Genomics, 100662), tissue was blocked at room temperature for 1 hour, stained overnight at 4C, and enhanced at room temperature for 20 minutes. Prepared tissue slides were loaded onto the Xenium Analyzer along with Xenium Decoding Consumables (10X Genomics, 100487) and Xenium Decoding Reagents (10X Genomics, 1000461) following the 10X Genomics protocol CG000584 Rev K for Xenium v1 Gene Expression Workflows.

### Analyses of HCMV gene expressions

#### TC Index Score

To order cells along a TC Index axis, we first calculated a module score for each temporal class group using the score_genes function from Scanpy (v1.11.5). For normalization, we then applied a softmax function to the resultant module scores to convert them to a probability distribution summing to 1 for every cell. Finally, we assigned an integer weight to each TC (e.g. TC1 = 1, TC2 = 2, TC3 = 3), and for each cell multiplied its softmax module score for each TC by that TC’s corresponding weight, then summed the weight to output a weighted progression score for each cell.

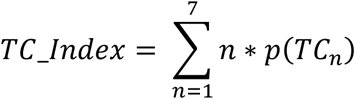

(Where *p*(*TC_n_*) *i*s the softmax probability associated with the module score for *TC_n_*) This TC Progression score corresponds to the transcriptional progression from expression of Early TC genes to Late TC genes, thus providing an estimate of progression through the lytic cycle.

#### TC Progression Classifier

To determine genes specific to low, medium, and high TC progression, we first ranked each cell by TC progression score, then split into 3 quantiles comprised of roughly equal cell number (n_low_ = 1999 cells, n_medium_ = 1998 cells, n_high_ = 1998 cells), that we labeled “low”, “medium”, and “high”. We then trained an eXtreme Gradient Boosting (XGBoost) classifier on the expression matrix (excluding HCMV viral genes) to predict each of the three quantile labels. The classifier model used the following parameters: n_estimators=300, max_depth=6, learning_rate=0.05, subsample=0.8, colsample_bytree=0.8, objective=’multi:softprob’, eval_metric=’mlogloss’, tree_method=’hist’. A validation set was generated by using an 80%-20% train-test split, stratified by quantile bin.

To unpack the model and analyze which genes contribute to differential classification of each quantile, we performed SHapley Additive exPlanations (SHAP) analysis on the test set (92). We plotted the genes contributing to the model on a beeswarm plot to illustrate top transcriptional contributions to the prediction of each class. We then conducted over-representation analysis (ORA) on the top 100 positive SHAP genes for each class (low, medium, and high) using Enrichr (93–95), and queried Gene Ontology (GO) Biological Processes 2023 (96,97), KEGG Pathway 2021 (98,99), MSigDB Hallmark 2020 (100), and Reactome 2022 (101). Results were filtered by False Discovery Rate (FDR) < 0.05.

#### Unsupervised Clustering

Unsupervised clustering was performed by Agglomerative Hierarchical Clustering using the Hierarchy module in SciPy 1.14.1. For linkage, we employed the Ward linkage method. The pairwise distance metric was computed by Pearson Correlation. To identify unique viral sub-states that represent samplings of different time points along infection progression, we isolated viral genes in the GFP+ subset of the HCMV-infected cortical organoid and blinded the clustering algorithm to host genes. We subsequently constructed a dendrogram to visualize clustering and select an optimal number of clusters (*k*). The cophenetic correlation coefficient (*c*) was calculated to determine preservation of pairwise distances within the structure of the dendrogram. Cophenetic correlation coefficient was found to be 0.4988, which we determined to be acceptable for the analyses given the primary focus on gaining a snapshot insight into different HCMV infection timepoints, rather than a discrete cluster annotation for each viral cluster. We then calculated the Silhouette coefficient across a range of *k*’s between *k=*2 and *k*=10. We chose several integer values for *k* (*k*=3, *k*=4, *k*=7) to sample different levels of coarseness in the unsupervised analysis of different infection timepoints. For the focus of downstream analyses, we chose to cut the dendrogram at *k*=7 (silhouette score = 0.1264) with the intention of balancing the visual structure of the dendrogram with a silhouette score acceptable for analysis of a semi-continuous progression through transcriptional timepoints during infection. To order the viral sub-state clusters along an infection progression axis, we calculated the average TC Progression score across all cells in each viral sub-state, then ordered each viral sub-state by increasing average TC Progression score. Significant difference in progression between viral sub-states was established using pairwise Mann-Whitney U and visualized using a violin plot.

To validate the utility of these clusters considering the modest silhouette scores, we trained an XGBoost classifier model to predict the cluster labels derived from hierarchical clustering, this time blinding the model to viral genes and including only host genes in the training data. Model parameters included the use of 500 estimators, a learning rate of 0.05, maximum tree depth of 6, row subsampling ratio of 0.8, and a column subsampling ratio of 0.8. The random state was initialized to 2. We then performed model validation using an 80%-20% train-test split, achieving a Receiver Operating Characteristic area under the curve (ROC-AUC) of 0.936. Finally, we constructed a confusion matrix to illustrate the prediction fidelity of our model.

### Differential Expression Analysis and Gene Set Enrichment Analysis (GSEA)

We performed differential expression analysis between each viral sub-state and all other cells using the rank_genes_groups function from Scanpy (v1.11.5). The top 10 differentially expressed genes for each viral sub-state were subsequently visualized by dot plot using the rank_genes_groups_dotplot function. To further analyze differentially expressed genes in each viral sub-state, we conducted pre-ranked Gene Set Enrichment Analysis (GSEA) on the entire set of differentially expressed genes sorted by log_2_ fold change using the GSEApy library (v1.1.13). We queried Gene Ontology (GO) Biological Processes 2023 (96,97), KEGG Pathway 2021 (98,99), MSigDB Hallmark 2020 (100), and Reactome 2022 (101). Results were filtered by False Discovery Rate (FDR) < 0.05.

### Spatial Transcriptomics

#### Data Preprocessing

Raw Xenium transcript data were re-segmented using ProSeg (v3.1.1) with the full transcript set to preserve spatial context across all tissue sections present on a given slide using the xenium preset. Output count matrices, cell metadata, and transcript metadata were used to construct objects using the Seurat (v5.4.0) in R for each organoid via CreateSeuratObject(). Cells were filtered to retain those with greater than 5 detected features and more than 10 total counts (nFeatureRNA > 5, nCount_RNA > 10). Since multiple organoids were embedded within the same capture area of each Xenium slide, organoids and their respective sections were separated based on their coordinates from the positioning image. Section boundaries were confirmed using histograms of the x-coordinate distribution and were determined individually for each organoid via visual inspection, as section positions and spacing varied across samples. Cells were then partitioned into discrete section-level Seurat objects based on manually defined x-coordinate cutoffs.

#### Robust Cell Type Deconvolution and downstream processing

To combat the limitation of spatial technology where individual measurements may contain contributions from multiple cells, thus limiting analysis of cell-type specific spatial patterns, and to integrate our spatial and scRNA-seq datasets, we utilized Robust Cell Type Decomposition (RCTD) using the R package spacexr (v2.2.0) (46,102). RCTD was run in doublet mode independently for each tissue section. Due to the lower per-cell transcript counts comparative to our scRNA-seq reference, UMI_min was lowered to 10, confidence_threshold was lowered to 2, and doublet_threshold was lowered to 8 from the default parameters (100, 5, and 20, respectively) following systematic testing of retention rates and classification quality across a range of values. Cells that could not be confidently assigned a cell type by RCTD were labeled as “unclassified” rather than assuming a specific biological state. To evaluate the accuracy of spatial deconvolution, we quantified the mean RCTD singlet assignment scores (singlets_score) across cell types in both HCMV-infected and mock organoids. Values were calculated for each cell population using singlet spots (spot_class == “singlet”). For cell populations exhibiting elevated singlet scores (e.g., Glutamatergic GABAergic Neurons), secondary cell-type predictions (second_type) and underlying spot-weight matrices were analyzed across all organoid biological replicates. Secondary-type frequency breakdowns and score distributions were quantified to distinguish between spatial multiplets and model misclassification.

Individual tissue sections were merged in two ways: per-organoid objects combining all sections from a single organoid, and condition-level objects combining all sections across all organoids within each experimental condition (HCMV and mock). Unclassified categories were excluded prior to spatial UMAP plotting. Spatial coordinate maps were generated using ggplot2 (v3.4.0) with fixed aspect ratios to preserve tissue orientation. Relative cell-type proportions were quantified for each organoid sample across experimental conditions. To determine whether cell types undergo spatial reorganization relative to the organoid core or periphery, normalized radial distance was calculated for each cell centroid within each organoid sample. To enable comparison across organoids of varying dimensions, distances were normalized relative to the maximum observed distance within each sample. To visualize this data, continuous spatial density profiles comparing HCMV and mock conditions were generated per cell type using smooth kernel density estimation (α=0.3). To statistically evaluate radial positioning shifts while accounting for organoid-level biological variation, Linear Mixed-Effects Models (LMMs) were fit for each cell type using the lme4 and lmerTest packages in R where experimental condition was modeled as a fixed effect and organoid sample identity was modeled as a random intercept.

### Immunofluorescence Staining

At 14 dpi, neurons and cortical organoids were fixed with 4% paraformaldehyde (PFA) overnight at 4°C. Coverslips with neurons were washed with phosphate-buffered saline (PBS) and stored until use. Cortical organoids were washed with PBS and placed in 30% sucrose in PBS. Fixed organoids were then embedded in a cryosectioning mold using optimum cutting temperature (OCT) compound (Fisher Scientific) and frozen on dry ice, then stored at −80°C. Cryosectioning was performed to obtain 10μm sections using a Leica cryostat. Sections were immediately mounted onto Fisherbrand Superfrost Plus Microscope Slides. Prior to immunofluorescence analysis, sections were allowed to dry for 1 hour and then washed 1 time with PBS.

Neuron coverslips and organoid sections were incubated in blocking buffer (5% normal donkey serum and 0.1% Triton-X 100 in PBS) for 1 hour, then incubated in primary antibody buffer (primary antibody, 5% normal donkey serum and 0.1% Triton-X 100 in PBS) overnight at 4°C. After primary incubation, samples were washed in PBS three times for 5 minutes and then incubated in secondary antibody buffer (secondary antibody, 5% normal donkey serum and 0.1% Triton-X 100 in PBS) for 1 hr at room temperature. Nuclei were stained using Hoechst 33342 (1:1,000 in PBS; Invitrogen) for 10 minutes. Following staining, Fluoromount-G Mounting Medium (Invitrogen) was applied over the samples, covered by glass coverslips and then sealed using clearcoat nail polish. Slides were imaged using a Zeis LSM980 confocal microscope, with image analysis conducted in Zeiss ZEN analysis software. Primary antibodies used were EPCAM (mouse, 14-9326-82, ThermoFisher) and TUJ1 (chicken, GTX85469; GeneTex). Species-appropriate fluorescent secondary antibodies were used.

### Data accessibility

Single cell RNAseq and spatial transcriptomic datasets have been deposited in the Gene Expression Omnibus (GEO) database under accession numbers GSE353289 and GSE352890, respectively. All other raw data are available upon reasonable request from the corresponding author.

## ACKNOWLEDGEMENTS

These studies were supported by the National Institute of Allergy and Infectious Diseases division of the National Institutes of Health under award numbers R01AI132414 and R01AI190029 (S.S.T. and A.D.E.) and through a philanthropic gift from the Stead Family Foundation (S.S.T). We would like to thank Jessica Gawrys, Donna McCallister, and the Malarkannan Lab for technical assistance with transcriptomic assays. We also would like to thank Drs. Rebekah Mokry and Suzette Rosas-Rogers for providing the viral stocks.

## Conceptualization

D.A.L., K.J.P., B.S.O., S.S.T., and A.D.E.; data curation: D.A.L., K.J.P., and B.S.O.; formal data analysis: D.A.L., K.J.P., B.S.O; C-W.L., K.T., S.S.T., and A.D.E.; funding acquisition and resources: S.S.T. and A.D.E.; writing of original draft, reviewing, and editing: D.A.L., K.J.P., S.S.T., and A.D.E.

## REFERENCES

1. Zuhair M, Smit GSA, Wallis G, Jabbar F, Smith C, Devleesschauwer B, et al. Estimation of the worldwide seroprevalence of cytomegalovirus: A systematic review and meta-analysis. Rev Med Virol. 2019 May;29(3):e2034. doi:10.1002/rmv.2034 PubMed PMID: 30706584.

2. Auriti C, Maddaloni C, Ronci S, Santisi A, Martini L, Dotta A, et al. Congenital Cytomegalovirus Infection in Pregnancy: Challenges in Early Diagnosis, Reinfection, and Secondary Prevention. Viruses. 2026 Jun 28;18(7):713. doi:10.3390/v18070713 PubMed PMID: 42515565; PubMed Central PMCID: PMC13431613.

3. Cheeran MCJ, Lokensgard JR, Schleiss MR. Neuropathogenesis of congenital cytomegalovirus infection: disease mechanisms and prospects for intervention. Clin Microbiol Rev. 2009 Jan;22(1):99–126, Table of Contents. doi:10.1128/CMR.00023-08 PubMed PMID: 19136436; PubMed Central PMCID: PMC2620634.

4. Chatzakis C, Shahar-Nissan K, Faure-Bardon V, Picone O, Hadar E, Amir J, et al. The effect of valacyclovir on secondary prevention of congenital cytomegalovirus infection, following primary maternal infection acquired periconceptionally or in the first trimester of pregnancy. An individual patient data meta-analysis. Am J Obstet Gynecol. 2024 Feb;230(2):109–117.e2. doi:10.1016/j.ajog.2023.07.022 PubMed PMID: 37473793.

5. Shahar-Nissan K, Pardo J, Peled O, Krause I, Bilavsky E, Wiznitzer A, et al. Valaciclovir to prevent vertical transmission of cytomegalovirus after maternal primary infection during pregnancy: a randomised, double-blind, placebo-controlled trial. Lancet. 2020 Sep 12;396(10253):779–85. doi:10.1016/S0140-6736(20)31868-7 PubMed PMID: 32919517.

6. Akingbola A, Adegbesan A, Adewole O, Adegoke K, Benson AE, Jombo PA, et al. The mRNA-1647 vaccine: A promising step toward the prevention of cytomegalovirus infection (CMV). Hum Vaccines Immunother. 2025 Dec;21(1):2450045. doi: 10.1080/21645515.2025.2450045 PubMed PMID: 39825496; PubMed Central PMCID: PMC12934182.

7. Chiaradia I, Lancaster MA. Brain organoids for the study of human neurobiology at the interface of in vitro and in vivo. Nat Neurosci. 2020 Dec;23(12):1496–508. doi:10.1038/s41593-020-00730-3 PubMed PMID: 33139941.

8. Saito A, Tankou S, Ishii K, Sakao-Suzuki M, Oh EC, Murdoch H, et al. Cytomegalovirus-encoded immediate early 1 protein perturbs neural progenitor proliferation via interfering with host PML-DISC1 interaction. J Biol Chem. 2026 Mar;302(3):111269. doi:10.1016/j.jbc.2026.111269 PubMed PMID: 41655694; PubMed Central PMCID: PMC12992096.

9. Huang SN, Pan YT, Zhou YP, Wang XZ, Mei MJ, Yang B, et al. Human Cytomegalovirus IE1 Impairs Neuronal Migration by Downregulating Connexin 43. J Virol. 2023 May 31;97(5):e0031323. doi:10.1128/jvi.00313-23 PubMed PMID: 37097169; PubMed Central PMCID: PMC10231247.

10. Wang XZ, Wen L, Zhou YP, Huang SN, Yang B, Cheng S, et al. Human cytomegalovirus pUL97 upregulates SOCS3 expression via transcription factor RFX7 in neural progenitor cells. PLoS Pathog. 2023 Feb;19(2):e1011166. doi:10.1371/journal.ppat.1011166 PubMed PMID: 36753521; PubMed Central PMCID: PMC9942973.

11. Adelman JW, Rosas-Rogers S, Schumacher ML, Mokry RL, Terhune SS, Ebert AD. Human cytomegalovirus induces significant structural and functional changes in terminally differentiated human cortical neurons. mBio. 2023 Dec 19;14(6):e0225123. doi:10.1128/mbio.02251-23 PubMed PMID: 37966250; PubMed Central PMCID: PMC10746155.

12. Adelman JW, Sukowaty AT, Partridge KJ, Gawrys JE, Akins A, Terhune SS, et al. HCMV infection of terminally differentiated neurons disrupts microtubule organization, resulting in neurite retraction. Microbiol Spectr. 2025 Nov 4;13(11):e0119825. doi:10.1128/spectrum.01198-25 PubMed PMID: 41059695; PubMed Central PMCID: PMC12584624.

13. He Z, Dony L, Fleck JS, Szałata A, Li KX, Slišković I, et al. An integrated transcriptomic cell atlas of human neural organoids. Nature. 2024 Nov;635(8039):690–8. doi:10.1038/s41586-024-08172-8 PubMed PMID: 39567792; PubMed Central PMCID: PMC11578878.

14. Mulder LA, Vieira da Sá R, Korsten J, Freeze E, Schotting AJ, Koen G, et al. Effect of antivirals on clinical and lab-adapted human cytomegalovirus strains using induced pluripotent stem cell-derived human neural models. Antiviral Res. 2025 Sep;241:106233. doi:10.1016/j.antiviral.2025.106233 PubMed PMID: 40618931.

15. Egilmezer E, Hamilton ST, Foster CSP, Marschall M, Rawlinson WD. Human cytomegalovirus (CMV) dysregulates neurodevelopmental pathways in cerebral organoids. Commun Biol. 2024 Mar 19;7(1):340. doi:10.1038/s42003-024-05923-1 PubMed PMID: 38504123; PubMed Central PMCID: PMC10951402.

16. O’Brien BS, Mokry RL, Schumacher ML, Rosas-Rogers S, Terhune SS, Ebert AD. Neutralizing antibodies with neurotropic factor treatment maintain neurodevelopmental gene expression upon exposure to human cytomegalovirus. J Virol. 2023 Oct 31;97(10):e0069623. doi:10.1128/jvi.00696-23 PubMed PMID: 37796129; PubMed Central PMCID: PMC10653813.

17. Ijezie EC, O’Dowd JM, Kuan MI, Faeth AR, Fortunato EA. HCMV Infection Reduces Nidogen-1 Expression, Contributing to Impaired Neural Rosette Development in Brain Organoids. J Virol. 2023 May 31;97(5):e0171822. doi:10.1128/jvi.01718-22 PubMed PMID: 37125912; PubMed Central PMCID: PMC10231252.

18. Mokry RL, O’Brien BS, Adelman JW, Rosas S, Schumacher ML, Ebert AD, et al. Nitric Oxide Attenuates Human Cytomegalovirus Infection yet Disrupts Neural Cell Differentiation and Tissue Organization. J Virol. 2022 Jul 27;96(14):e0012622. doi:10.1128/jvi.00126-22 PubMed PMID: 35862705; PubMed Central PMCID: PMC9327702.

19. O’Brien BS, Mokry RL, Schumacher ML, Pulakanti K, Rao S, Terhune SS, et al. Downregulation of neurodevelopmental gene expression in iPSC-derived cerebral organoids upon infection by human cytomegalovirus. iScience. 2022 Apr 15;25(4):104098. doi:10.1016/j.isci.2022.104098 PubMed PMID: 35391828; PubMed Central PMCID: PMC8980761.

20. Sun G, Chiuppesi F, Chen X, Wang C, Tian E, Nguyen J, et al. Modeling Human Cytomegalovirus-Induced Microcephaly in Human iPSC-Derived Brain Organoids. Cell Rep Med. 2020 Apr 21;1(1):100002. doi:10.1016/j.xcrm.2020.100002 PubMed PMID: 33205055; PubMed Central PMCID: PMC7659592.

21. Sison SL, O’Brien BS, Johnson AJ, Seminary ER, Terhune SS, Ebert AD. Human Cytomegalovirus Disruption of Calcium Signaling in Neural Progenitor Cells and Organoids. J Virol. 2019 Sep 1;93(17):e00954–19. doi:10.1128/JVI.00954-19 PubMed PMID: 31217241; PubMed Central PMCID: PMC6694809.

22. Brown RM, Rana PSJB, Jaeger HK, O’Dowd JM, Balemba OB, Fortunato EA. Human Cytomegalovirus Compromises Development of Cerebral Organoids. J Virol. 2019 Sep 1;93(17):e00957–19. doi:10.1128/JVI.00957-19 PubMed PMID: 31217239; PubMed Central PMCID: PMC6694831.

23. Liu XJ, Jiang X, Huang SN, Sun JY, Zhao F, Zeng WB, et al. Human cytomegalovirus infection dysregulates neural progenitor cell fate by disrupting Hes1 rhythm and down-regulating its expression. Virol Sin. 2017 Jun;32(3):188–98. doi:10.1007/s12250-017-3956-0 PubMed PMID: 28451898; PubMed Central PMCID: PMC6598910.

24. Liu XJ, Yang B, Huang SN, Wu CC, Li XJ, Cheng S, et al. Human cytomegalovirus IE1 downregulates Hes1 in neural progenitor cells as a potential E3 ubiquitin ligase. PLoS Pathog. 2017 Jul;13(7):e1006542. doi:10.1371/journal.ppat.1006542 PubMed PMID: 28750047; PubMed Central PMCID: PMC5549770.

25. Li XJ, Liu XJ, Yang B, Fu YR, Zhao F, Shen ZZ, et al. Human Cytomegalovirus Infection Dysregulates the Localization and Stability of NICD1 and Jag1 in Neural Progenitor Cells. J Virol. 2015 Jul;89(13):6792–804. doi:10.1128/JVI.00351-15 PubMed PMID: 25903338; PubMed Central PMCID: PMC4468470.

26. Luo MH, Hannemann H, Kulkarni AS, Schwartz PH, O’Dowd JM, Fortunato EA. Human cytomegalovirus infection causes premature and abnormal differentiation of human neural progenitor cells. J Virol. 2010 Apr;84(7):3528–41. doi:10.1128/JVI.02161-09 PubMed PMID: 20071566; PubMed Central PMCID: PMC2838134.

27. Pan X, Li XJ, Liu XJ, Yuan H, Li JF, Duan YL, et al. Later passages of neural progenitor cells from neonatal brain are more permissive for human cytomegalovirus infection. J Virol. 2013 Oct;87(20):10968–79. doi:10.1128/JVI.01120-13 PubMed PMID: 23903847; PubMed Central PMCID: PMC3807278.

28. Nakamura H, Liao H, Minami K, Toyoda M, Akutsu H, Miyagawa Y, et al. Human cytomegalovirus induces apoptosis in neural stem/progenitor cells derived from induced pluripotent stem cells by generating mitochondrial dysfunction and endoplasmic reticulum stress. Herpesviridae. 2013 Oct 21;4(1):2. doi:10.1186/2042-4280-4-2 PubMed PMID: 24144363; PubMed Central PMCID: PMC3875896.

29. Wu CC, Jiang X, Wang XZ, Liu XJ, Li XJ, Yang B, et al. Human Cytomegalovirus Immediate Early 1 Protein Causes Loss of SOX2 from Neural Progenitor Cells by Trapping Unphosphorylated STAT3 in the Nucleus. J Virol. 2018 Sep 1;92(17):e00340–18. doi:10.1128/JVI.00340-18 PubMed PMID: 29950413; PubMed Central PMCID: PMC6096794.

30. D’Aiuto L, Di Maio R, Heath B, Raimondi G, Milosevic J, Watson AM, et al. Human induced pluripotent stem cell-derived models to investigate human cytomegalovirus infection in neural cells. PloS One. 2012;7(11):e49700. doi:10.1371/journal.pone.0049700 PubMed PMID: 23209593; PubMed Central PMCID: PMC3507916.

31. Mercorelli B, Poli E, Pianezzola A, Faggin E, Arav-Boger R, Palù G, et al. Letermovir shows antiviral and neuroprotective effects in differentiating neurons and cerebral organoids mimicking human developing brain. Cell Mol Life Sci CMLS. 2026 Jan 11;83(1):139. doi:10.1007/s00018-025-06015-x PubMed PMID: 41520074; PubMed Central PMCID: PMC12979744.

32. van den Pol AN, Robek MD, Ghosh PK, Ozduman K, Bandi P, Whim MD, et al. Cytomegalovirus induces interferon-stimulated gene expression and is attenuated by interferon in the developing brain. J Virol. 2007 Jan;81(1):332–48. doi:10.1128/JVI.01592-06 PubMed PMID: 17065212; PubMed Central PMCID: PMC1797251.

33. Goodrum F, Britt W, Mocarski ES. Cytomegalovirus. In: Fields Virology: DNA Viruses. 7th ed. Philadelphia: Wolters Kluwer; 2022. p. 389–444.

34. Rozman B, Nachshon A, Levi Samia R, Lavi M, Schwartz M, Stern-Ginossar N. Temporal dynamics of HCMV gene expression in lytic and latent infections. Cell Rep. 2022 Apr 12;39(2):110653. doi:10.1016/j.celrep.2022.110653 PubMed PMID: 35417700; PubMed Central PMCID: PMC9035752.

35. Chambers J, Angulo A, Amaratunga D, Guo H, Jiang Y, Wan JS, et al. DNA microarrays of the complex human cytomegalovirus genome: profiling kinetic class with drug sensitivity of viral gene expression. J Virol. 1999 Jul;73(7):5757–66. doi:10.1128/JVI.73.7.5757-5766.1999 PubMed PMID: 10364327; PubMed Central PMCID: PMC112636.

36. Yaiw KC, Ferrer IA, Söderberg-Nauclér C, Religa D. Human cytomegalovirus infection and cognitive decline: insights from population and experimental studies. Front Aging Neurosci. 2025;17:1757461. doi:10.3389/fnagi.2025.1757461 PubMed PMID: 41641002; PubMed Central PMCID: PMC12865707.

37. Sanami S, Shamsabadi S, Dayhimi A, Pirhayati M, Ahmad S, Pirhayati A, et al. Association between cytomegalovirus infection and neurological disorders: A systematic review. Rev Med Virol. 2024 May;34(3):e2532. doi:10.1002/rmv.2532 PubMed PMID: 38549138.

38. Lancaster MA, Knoblich JA. Generation of cerebral organoids from human pluripotent stem cells. Nat Protoc. 2014 Oct;9(10):2329–40. doi:10.1038/nprot.2014.158 PubMed PMID: 25188634; PubMed Central PMCID: PMC4160653.

39. Sinzger C, Hahn G, Digel M, Katona R, Sampaio KL, Messerle M, et al. Cloning and sequencing of a highly productive, endotheliotropic virus strain derived from human cytomegalovirus TB40/E. J Gen Virol. 2008 Feb;89(Pt 2):359–68. doi:10.1099/vir.0.83286-0 PubMed PMID: 18198366.

40. He Z, Dony L, Fleck JS, Szałata A, Li KX, Slišković I, et al. An integrated transcriptomic cell atlas of human neural organoids. Nature. 2024 Nov 21;635(8039):690–8. doi:10.1038/s41586-024-08172-8

41. Velasco S, Kedaigle AJ, Simmons SK, Nash A, Rocha M, Quadrato G, et al. Individual brain organoids reproducibly form cell diversity of the human cerebral cortex. Nature. 2019 Jun;570(7762):523–7. doi:10.1038/s41586-019-1289-x

42. Crowley G, Tabula Sapiens Consortium, Jones RC, Krasnow M, Pisco AO, Salzman J, et al. Benchmarking cell type and gene set annotation by large language models with AnnDictionary. Nat Commun. 2025 Oct 28;16(1):9511. doi:10.1038/s41467-025-64511-x

43. Anthropic. Claude Haiku 4.5 [Internet]. San Francisco (CA): Anthropic; 2025. (claude-haiku-4-5). Available from: https://docs.claude.com

44. www.atlasantibodies.com [Internet]. [cited 2026 Jul 7]. Cortical Layer Markers. Available from: https://www.atlasantibodies.com/knowledge-hub/white-papers/cortical-layer-markers/?language=en_GB

45. Rozman B, Nachshon A, Levi Samia R, Lavi M, Schwartz M, Stern-Ginossar N. Temporal dynamics of HCMV gene expression in lytic and latent infections. Cell Rep. 2022 Apr;39(2):110653. doi:10.1016/j.celrep.2022.110653

46. Cable DM, Murray E, Zou LS, Goeva A, Macosko EZ, Chen F, et al. Robust decomposition of cell type mixtures in spatial transcriptomics. Nat Biotechnol. 2022 Apr;40(4):517–26. doi:10.1038/s41587-021-00830-w PubMed PMID: 33603203; PubMed Central PMCID: PMC8606190.

47. Demetci P, Santorella R, Sandstede B, Noble WS, Singh R. SCOT: Single-Cell Multi-Omics Alignment with Optimal Transport. J Comput Biol. 2022 Jan 1;29(1):3–18. doi:10.1089/cmb.2021.0446

48. Klein D, Palla G, Lange M, Klein M, Piran Z, Gander M, et al. Mapping cells through time and space with moscot. Nature. 2025 Feb 27;638(8052):1065–75. doi:10.1038/s41586-024-08453-2

49. Schiebinger G, Shu J, Tabaka M, Cleary B, Subramanian V, Solomon A, et al. Optimal-Transport Analysis of Single-Cell Gene Expression Identifies Developmental Trajectories in Reprogramming. Cell. 2019 Feb;176(4):928–943.e22. doi:10.1016/j.cell.2019.01.006

50. Gerna G, Kabanova A, Lilleri D. Human Cytomegalovirus Cell Tropism and Host Cell Receptors. Vaccines. 2019 Jul 22;7(3):70. doi:10.3390/vaccines7030070 PubMed PMID: 31336680; PubMed Central PMCID: PMC6789482.

51. Ward JH. Hierarchical Grouping to Optimize an Objective Function. J Am Stat Assoc. 1963 Mar;58(301):236–44. doi:10.1080/01621459.1963.10500845

52. Kalluri R, Weinberg RA. The basics of epithelial-mesenchymal transition. J Clin Invest. 2009 Jun;119(6):1420–8. doi:10.1172/JCI39104 PubMed PMID: 19487818; PubMed Central PMCID: PMC2689101.

53. Mahmud J, Nandi I, Procter DJ, Walsh D. Cytomegalovirus disrupts Lamin A/C to control microtubule-mediated nuclear movement and cell migration. Proc Natl Acad Sci U S A. 2025 Dec 2;122(48):e2507831122. doi:10.1073/pnas.2507831122 PubMed PMID: 41289383; PubMed Central PMCID: PMC12685118.

54. Luganini A, Serra V, Scarpellino G, Bhat SM, Munaron L, Fiorio Pla A, et al. The US21 viroporin of human cytomegalovirus stimulates cell migration and adhesion. mBio. 2023 Aug 31;14(4):e0074923. doi:10.1128/mbio.00749-23 PubMed PMID: 37477430; PubMed Central PMCID: PMC10470750.

55. Lau B, Kerr K, Camiolo S, Nightingale K, Gu Q, Antrobus R, et al. Human Cytomegalovirus RNA2.7 Is Required for Upregulating Multiple Cellular Genes To Promote Cell Motility and Viral Spread Late in Lytic Infection. J Virol. 2021 Sep 27;95(20):e0069821. doi:10.1128/JVI.00698-21 PubMed PMID: 34346763; PubMed Central PMCID: PMC8475523.

56. Streblow DN, Soderberg-Naucler C, Vieira J, Smith P, Wakabayashi E, Ruchti F, et al. The human cytomegalovirus chemokine receptor US28 mediates vascular smooth muscle cell migration. Cell. 1999 Nov 24;99(5):511–20. doi:10.1016/s0092-8674(00)81539-1 PubMed PMID: 10589679.

57. Galaburda AM. Developmental dyslexia and animal studies: at the interface between cognition and neurology. Cognition. 1994;50(1–3):133–49. doi:10.1016/0010-0277(94)90025-6 PubMed PMID: 8039357.

58. Ghari M, Ghaemimanesh F. EpCAM: A multifunctional protein at the crossroads of cancer biology, diagnosis, and immunotherapy. Cancer Treat Res Commun. 2026;48:101258. doi:10.1016/j.ctarc.2026.101258 PubMed PMID: 42235112.

59. Olbrich H, Theobald SJ, Slabik C, Gerasch L, Schneider A, Mach M, et al. Adult and Cord Blood-Derived High-Affinity gB-CAR-T Cells Effectively React Against Human Cytomegalovirus Infections. Hum Gene Ther. 2020 Apr;31(7–8):423–39. doi:10.1089/hum.2019.149 PubMed PMID: 32159399; PubMed Central PMCID: PMC7194322.

60. Ali A, Chiuppesi F, Nguyen M, Hausner MA, Nguyen J, Kha M, et al. Chimeric Antigen Receptors Targeting Human Cytomegalovirus. J Infect Dis. 2020 Aug 4;222(5):853–62. doi:10.1093/infdis/jiaa171 PubMed PMID: 32285133; PubMed Central PMCID: PMC7399702.

61. Keller MD, Hanley PJ, Chi YY, Aguayo-Hiraldo P, Dvorak CC, Verneris MR, et al. Antiviral cellular therapy for enhancing T-cell reconstitution before or after hematopoietic stem cell transplantation (ACES): a two-arm, open label phase II interventional trial of pediatric patients with risk factor assessment. Nat Commun. 2024 Apr 18;15(1):3258. doi:10.1038/s41467-024-47057-2 PubMed PMID: 38637498; PubMed Central PMCID: PMC11026387.

62. Shimamura M, Murphy-Ullrich JE, Britt WJ. Human cytomegalovirus induces TGF-β1 activation in renal tubular epithelial cells after epithelial-to-mesenchymal transition. PLoS Pathog. 2010 Nov 4;6(11):e1001170. doi:10.1371/journal.ppat.1001170 PubMed PMID: 21079788; PubMed Central PMCID: PMC2973835.

63. Ye Z, Hu X, Rahaman SM, Zheng Y, Zhang X, Tang CSM, et al. HCMV infection disrupts barrier functions and promotes epithelial-mesenchymal transition in a cholangiocyte organoid model. Nat Commun. 2026 Feb 3;17(1):2273. doi:10.1038/s41467-026-68962-8 PubMed PMID: 41634045; PubMed Central PMCID: PMC12966431.

64. Yoo YD, Chiou CJ, Choi KS, Yi Y, Michelson S, Kim S, et al. The IE2 regulatory protein of human cytomegalovirus induces expression of the human transforming growth factor beta1 gene through an Egr-1 binding site. J Virol. 1996 Oct;70(10):7062–70. doi:10.1128/JVI.70.10.7062-7070.1996 PubMed PMID: 8794351; PubMed Central PMCID: PMC190757.

65. Michelson S, Alcami J, Kim SJ, Danielpour D, Bachelerie F, Picard L, et al. Human cytomegalovirus infection induces transcription and secretion of transforming growth factor beta 1. J Virol. 1994 Sep;68(9):5730–7. doi:10.1128/JVI.68.9.5730-5737.1994 PubMed PMID: 8057454; PubMed Central PMCID: PMC236976.

66. Golconda P, Andrade-Medina M, Oberstein A. Subconfluent ARPE-19 Cells Display Mesenchymal Cell-State Characteristics and Behave like Fibroblasts, Rather Than Epithelial Cells, in Experimental HCMV Infection Studies. Viruses. 2023 Dec 28;16(1):49. doi:10.3390/v16010049 PubMed PMID: 38257749; PubMed Central PMCID: PMC10821009.

67. Oberstein A, Shenk T. Cellular responses to human cytomegalovirus infection: Induction of a mesenchymal-to-epithelial transition (MET) phenotype. Proc Natl Acad Sci U S A. 2017 Sep 26;114(39):E8244–53. doi:10.1073/pnas.1710799114 PubMed PMID: 28874566; PubMed Central PMCID: PMC5625929.

68. Moreno I, Rodríguez-Sánchez I, Schafer X, Munger J. Human cytomegalovirus induces neuronal enolase to support virally mediated metabolic remodeling. Proc Natl Acad Sci U S A. 2022 Dec 6;119(49):e2205789119. doi:10.1073/pnas.2205789119 PubMed PMID: 36459650; PubMed Central PMCID: PMC9894225.

69. Kelnhofer-Millevolte LE, Smith JR, Nguyen DH, Wilson LS, Lewis HC, Arnold EA, et al. Human cytomegalovirus induces neuronal gene expression through IE1 for viral maturation. Nat Commun. 2025 Aug 8;16(1):7316. doi:10.1038/s41467-025-61915-7 PubMed PMID: 40781068; PubMed Central PMCID: PMC12334593.

70. González P, Tuñón MJ, Manrique V, Garcia-Pardo LA, González J. Changes in hepatic cytosolic glutathione S-transferase enzymes induced by clotrimazole treatment in rats. Clin Exp Pharmacol Physiol. 1989 Nov;16(11):867–71. doi:10.1111/j.1440-1681.1989.tb01526.x PubMed PMID: 2612064.

71. Bogdanow B, Phan QV, Wiebusch L. Emerging Mechanisms of G1/S Cell Cycle Control by Human and Mouse Cytomegaloviruses. mBio. 2021 Dec 21;12(6):e0293421. doi:10.1128/mBio.02934-21 PubMed PMID: 34903047; PubMed Central PMCID: PMC8669474.

72. Hagemeier C, Caswell R, Hayhurst G, Sinclair J, Kouzarides T. Functional interaction between the HCMV IE2 transactivator and the retinoblastoma protein. EMBO J. 1994 Jun 15;13(12):2897–903. doi:10.1002/j.1460-2075.1994.tb06584.x PubMed PMID: 8026474; PubMed Central PMCID: PMC395171.

73. Iwahori S, Umaña AC, VanDeusen HR, Kalejta RF. Human cytomegalovirus-encoded viral cyclin-dependent kinase (v-CDK) UL97 phosphorylates and inactivates the retinoblastoma protein-related p107 and p130 proteins. J Biol Chem. 2017 Apr 21;292(16):6583–99. doi:10.1074/jbc.M116.773150 PubMed PMID: 28289097; PubMed Central PMCID: PMC5399109.

74. Fortunato EA, Sommer MH, Yoder K, Spector DH. Identification of domains within the human cytomegalovirus major immediate-early 86-kilodalton protein and the retinoblastoma protein required for physical and functional interaction with each other. J Virol. 1997 Nov;71(11):8176–85. doi:10.1128/JVI.71.11.8176-8185.1997 PubMed PMID: 9343168; PubMed Central PMCID: PMC192274.

75. Iwahori S, Hakki M, Chou S, Kalejta RF. Molecular Determinants for the Inactivation of the Retinoblastoma Tumor Suppressor by the Viral Cyclin-dependent Kinase UL97. J Biol Chem. 2015 Aug 7;290(32):19666–80. doi:10.1074/jbc.M115.660043 PubMed PMID: 26100623; PubMed Central PMCID: PMC4528131.

76. Hume AJ, Finkel JS, Kamil JP, Coen DM, Culbertson MR, Kalejta RF. Phosphorylation of retinoblastoma protein by viral protein with cyclin-dependent kinase function. Science. 2008 May 9;320(5877):797–9. doi:10.1126/science.1152095 PubMed PMID: 18467589.

77. Iwahori S, Kalejta RF. Phosphorylation of transcriptional regulators in the retinoblastoma protein pathway by UL97, the viral cyclin-dependent kinase encoded by human cytomegalovirus. Virology. 2017 Dec;512:95–103. doi:10.1016/j.virol.2017.09.009 PubMed PMID: 28946006; PubMed Central PMCID: PMC5653418.

78. Rodríguez-Sánchez I, Schafer XL, Monaghan M, Munger J. The Human Cytomegalovirus UL38 protein drives mTOR-independent metabolic flux reprogramming by inhibiting TSC2. PLoS Pathog. 2019 Jan;15(1):e1007569. doi:10.1371/journal.ppat.1007569 PubMed PMID: 30677091; PubMed Central PMCID: PMC6363234.

79. Bai Y, Xuan B, Liu H, Zhong J, Yu D, Qian Z. Tuberous Sclerosis Complex Protein 2-Independent Activation of mTORC1 by Human Cytomegalovirus pUL38. J Virol. 2015 Aug;89(15):7625–35. doi:10.1128/JVI.01027-15 PubMed PMID: 25972538; PubMed Central PMCID: PMC4505643.

80. Qian Z, Xuan B, Gualberto N, Yu D. The human cytomegalovirus protein pUL38 suppresses endoplasmic reticulum stress-mediated cell death independently of its ability to induce mTORC1 activation. J Virol. 2011 Sep;85(17):9103–13. doi:10.1128/JVI.00572-11 PubMed PMID: 21715486; PubMed Central PMCID: PMC3165799.

81. Moorman NJ, Cristea IM, Terhune SS, Rout MP, Chait BT, Shenk T. Human cytomegalovirus protein UL38 inhibits host cell stress responses by antagonizing the tuberous sclerosis protein complex. Cell Host Microbe. 2008 Apr 17;3(4):253–62. doi:10.1016/j.chom.2008.03.002 PubMed PMID: 18407068; PubMed Central PMCID: PMC2759192.

82. Yu Y, Maguire TG, Alwine JC. Human cytomegalovirus infection induces adipocyte-like lipogenesis through activation of sterol regulatory element binding protein 1. J Virol. 2012 Mar;86(6):2942–9. doi:10.1128/JVI.06467-11 PubMed PMID: 22258239; PubMed Central PMCID: PMC3302344.

83. Yu Y, Pierciey FJ, Maguire TG, Alwine JC. PKR-like endoplasmic reticulum kinase is necessary for lipogenic activation during HCMV infection. PLoS Pathog. 2013;9(4):e1003266. doi:10.1371/journal.ppat.1003266 PubMed PMID: 23592989; PubMed Central PMCID: PMC3617203.

84. Seo JY, Cresswell P. Viperin regulates cellular lipid metabolism during human cytomegalovirus infection. PLoS Pathog. 2013;9(8):e1003497. doi:10.1371/journal.ppat.1003497 PubMed PMID: 23935494; PubMed Central PMCID: PMC3731232.

85. Yu Y, Maguire TG, Alwine JC. ChREBP, a glucose-responsive transcriptional factor, enhances glucose metabolism to support biosynthesis in human cytomegalovirus-infected cells. Proc Natl Acad Sci U S A. 2014 Feb 4;111(5):1951–6. doi:10.1073/pnas.1310779111 PubMed PMID: 24449882; PubMed Central PMCID: PMC3918764.

86. Vysochan A, Sengupta A, Weljie AM, Alwine JC, Yu Y. ACSS2-mediated acetyl-CoA synthesis from acetate is necessary for human cytomegalovirus infection. Proc Natl Acad Sci U S A. 2017 Feb 21;114(8):E1528–35. doi:10.1073/pnas.1614268114 PubMed PMID: 28167750; PubMed Central PMCID: PMC5338361.

87. Raymonda MH, Rodríguez-Sánchez I, Schafer XL, Smorodintsev-Schiller L, Harris IS, Munger J. Cytomegalovirus-induced inactivation of TSC2 disrupts the coupling of fatty acid biosynthesis to glucose availability resulting in a vulnerability to glucose starvation. mBio. 2024 Jan 16;15(1):e0303123. doi:10.1128/mbio.03031-23 PubMed PMID: 38117060; PubMed Central PMCID: PMC10790783.

88. Mokry RL, de Lacy CB, Purdy JG. Glutamine supports human cytomegalovirus induced lipidome remodeling through reductive carboxylation. PLoS Pathog. 2026 Aug;22(8):e1014496. doi:10.1371/journal.ppat.1014496 PubMed PMID: 42585263; PubMed Central PMCID: PMC13489524.

89. Isler JA, Skalet AH, Alwine JC. Human cytomegalovirus infection activates and regulates the unfolded protein response. J Virol. 2005 Jun;79(11):6890–9. doi:10.1128/JVI.79.11.6890-6899.2005 PubMed PMID: 15890928; PubMed Central PMCID: PMC1112127.

90. Siddiquey MNA, Zhang H, Nguyen CC, Domma AJ, Kamil JP. The Human Cytomegalovirus Endoplasmic Reticulum-Resident Glycoprotein UL148 Activates the Unfolded Protein Response. J Virol. 2018 Oct 15;92(20):e00896–18. doi:10.1128/JVI.00896-18 PubMed PMID: 30045994; PubMed Central PMCID: PMC6158444.

91. Zhang H, Read C, Nguyen CC, Siddiquey MNA, Shang C, Hall CM, et al. The Human Cytomegalovirus Nonstructural Glycoprotein UL148 Reorganizes the Endoplasmic Reticulum. mBio. 2019 Dec 10;10(6):e02110–19. doi:10.1128/mBio.02110-19 PubMed PMID: 31822584; PubMed Central PMCID: PMC6904874.

92. Lundberg SM, Lee SI. A Unified Approach to Interpreting Model Predictions.

93. Kuleshov MV, Jones MR, Rouillard AD, Fernandez NF, Duan Q, Wang Z, et al. Enrichr: a comprehensive gene set enrichment analysis web server 2016 update. Nucleic Acids Res. 2016 Jul 8;44(W1):W90–97. doi:10.1093/nar/gkw377 PubMed PMID: 27141961; PubMed Central PMCID: PMC4987924.

94. Chen EY, Tan CM, Kou Y, Duan Q, Wang Z, Meirelles GV, et al. Enrichr: interactive and collaborative HTML5 gene list enrichment analysis tool. BMC Bioinformatics. 2013 Apr 15;14:128. doi:10.1186/1471-2105-14-128 PubMed PMID: 23586463; PubMed Central PMCID: PMC3637064.

95. Xie Z, Bailey A, Kuleshov MV, Clarke DJB, Evangelista JE, Jenkins SL, et al. Gene Set Knowledge Discovery with Enrichr. Curr Protoc. 2021 Mar;1(3):e90. doi:10.1002/cpz1.90

96. Ashburner M, Ball CA, Blake JA, Botstein D, Butler H, Cherry JM, et al. Gene Ontology: tool for the unification of biology. Nat Genet. 2000 May;25(1):25–9. doi:10.1038/75556

97. The Gene Ontology Consortium, Aleksander SA, Balhoff J, Carbon S, Cherry JM, Drabkin HJ, et al. The Gene Ontology knowledgebase in 2023. Baryshnikova A, editor. GENETICS. 2023 May 4;224(1):iyad031. doi:10.1093/genetics/iyad031

98. Kanehisa M, Furumichi M, Sato Y, Ishiguro-Watanabe M, Tanabe M. KEGG: integrating viruses and cellular organisms. Nucleic Acids Res. 2021 Jan 8;49(D1):D545–51. doi:10.1093/nar/gkaa970

99. Kanehisa M. KEGG: Kyoto Encyclopedia of Genes and Genomes. Nucleic Acids Res. 2000 Jan 1;28(1):27–30. doi:10.1093/nar/28.1.27

100. Liberzon A, Birger C, Thorvaldsdóttir H, Ghandi M, Mesirov JP, Tamayo P. The Molecular Signatures Database Hallmark Gene Set Collection. Cell Syst. 2015 Dec;1(6):417–25. doi:10.1016/j.cels.2015.12.004

101. Gillespie M, Jassal B, Stephan R, Milacic M, Rothfels K, Senff-Ribeiro A, et al. The reactome pathway knowledgebase 2022. Nucleic Acids Res. 2022 Jan 7;50(D1):D687–92. doi:10.1093/nar/gkab1028

102. Cable DM, Murray E, Shanmugam V, Zhang S, Zou LS, Diao M, et al. Cell type-specific inference of differential expression in spatial transcriptomics. Nat Methods. 2022 Sep;19(9):1076–87. doi:10.1038/s41592-022-01575-3 PubMed PMID: 36050488; PubMed Central PMCID: PMC10463137.

