## Supporting Information for "Cytomegalovirus Infection in Human Cortical Organoids Induces Unique Developmental and Viral Substates"

**SUPPORTING INFORMATION (Laws et al.)**

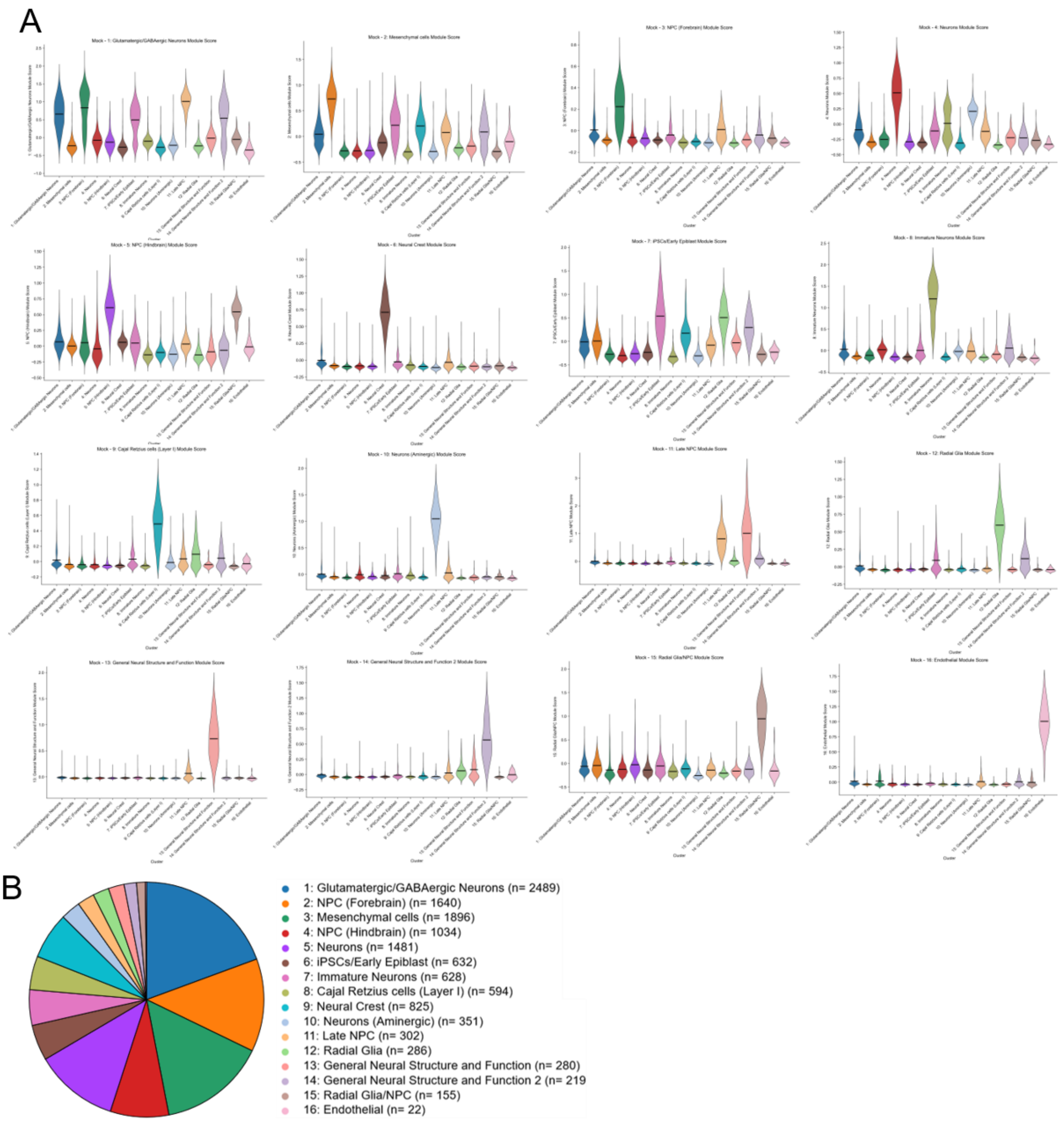


**S1 Fig. Uniqueness of cell type identity and cell type composition analysis in Mock cortical organoid.** (A) Violin Plot of Scanpy module scores of each of the sets of marker genes used to define each cluster. Significantly elevated module scores illustrate the uniqueness of the gene set used to annotate each cluster. (B) Circle chart showing cell type composition of the Mock cortical organoid.

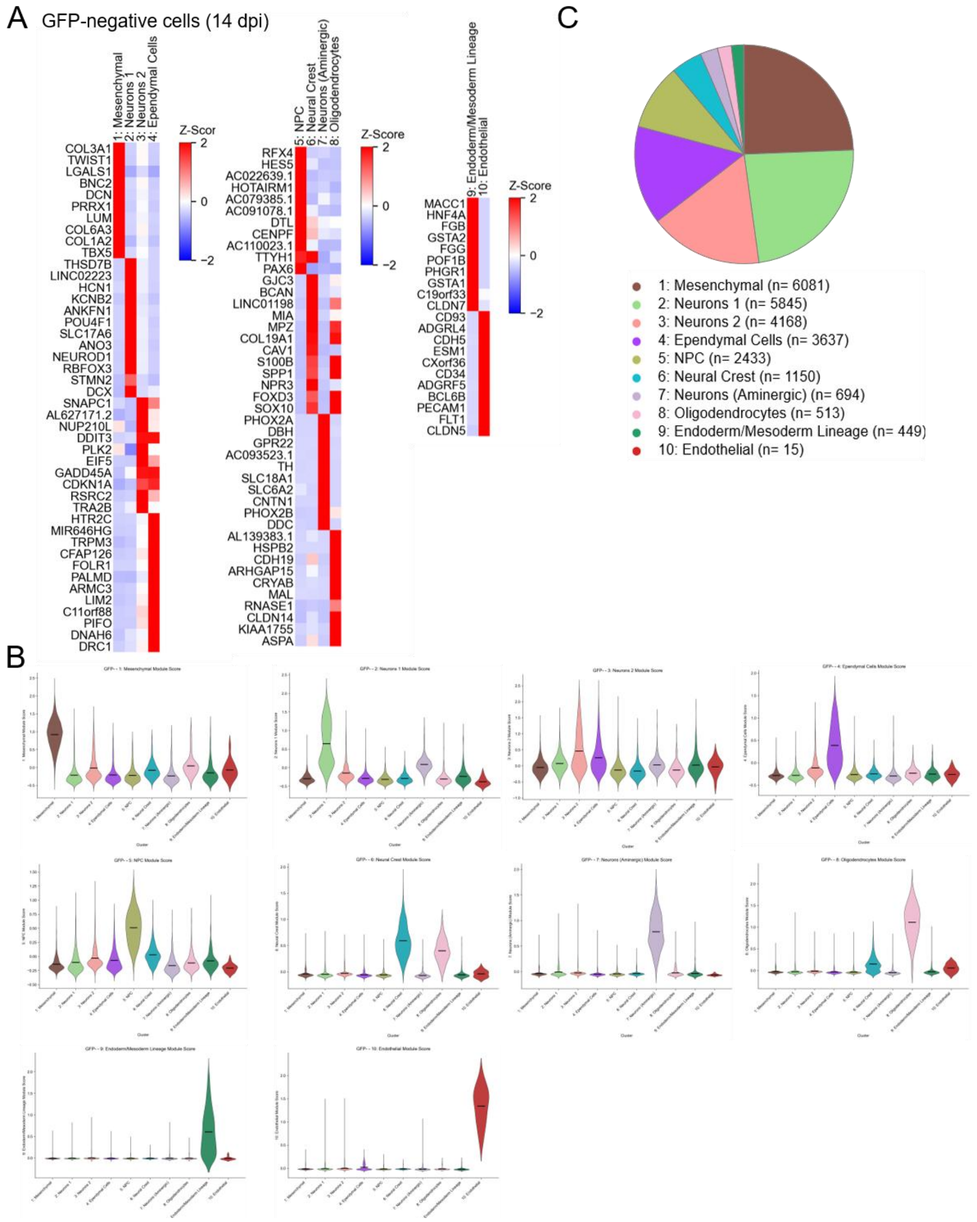


**S2 Fig. Cell type annotation markers and cell type composition of GFP-negative subset of the HCMV-infected cortical organoid.** (A) Heatmap of top marker genes for each cell type cluster for the GFP-negative subset of the HCMV infected organoid. (B) Violin plot of module scores for the top marker genes of each cell type cluster in the GFP- subset of the HCMV infected organoid, illustrating that each cluster uniquely expresses its own markers. (C) Circle chart showing cell type composition of the GFP-negative subset of the HCMV-infected cortical organoid.

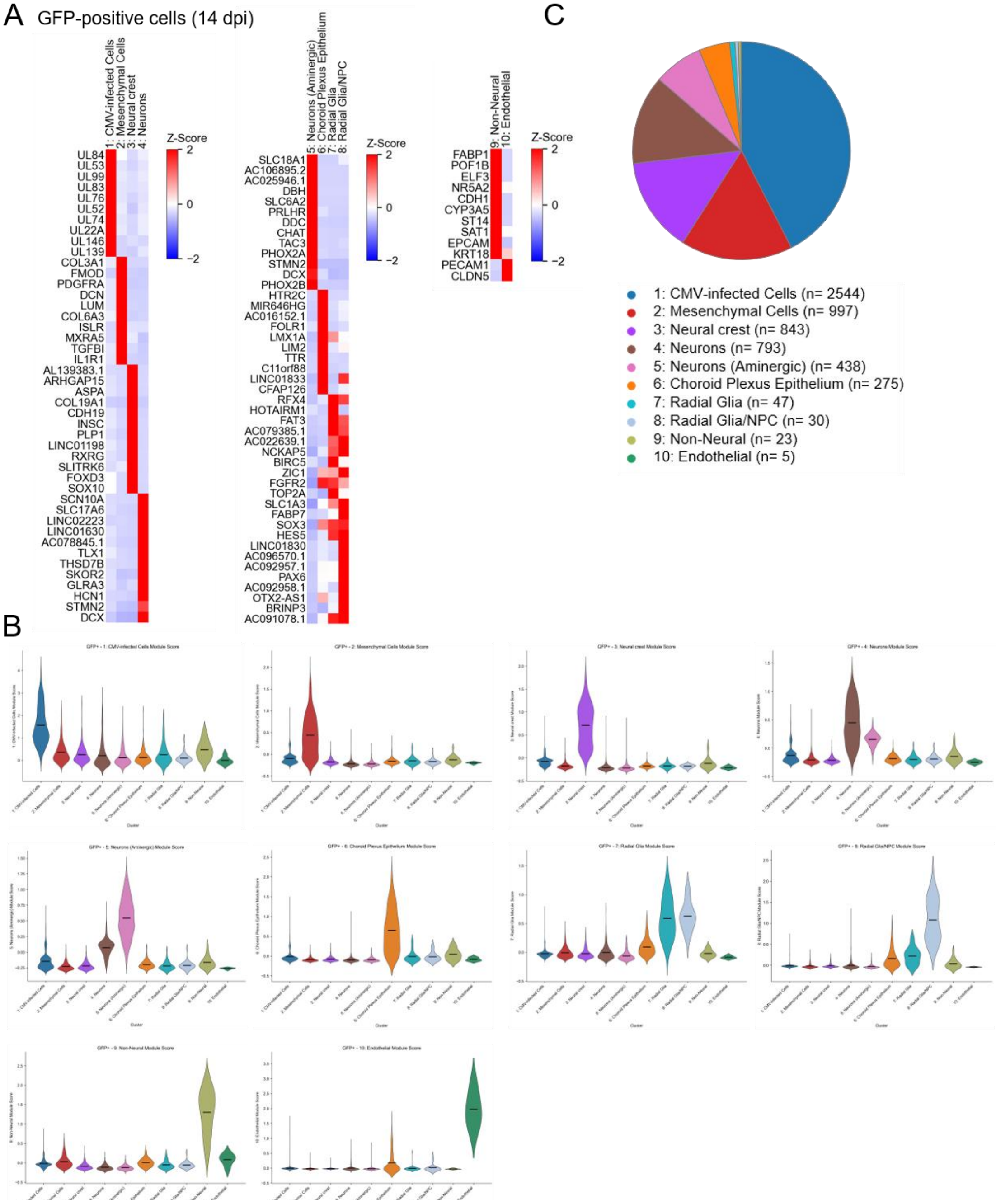


**S3 Fig. Cell type annotation markers and cell type composition of GFP-positive subset of the HCMV-infected cortical organoid.** (A) Heatmap showing cell type marker expression across cell types in the GFP-positive subset of the HCMV-infected organoid. (B) Module scores of top marker genes for each cell type cluster in GFP-positive. (C) Circle chart of cell type composition of the GFP-positive.

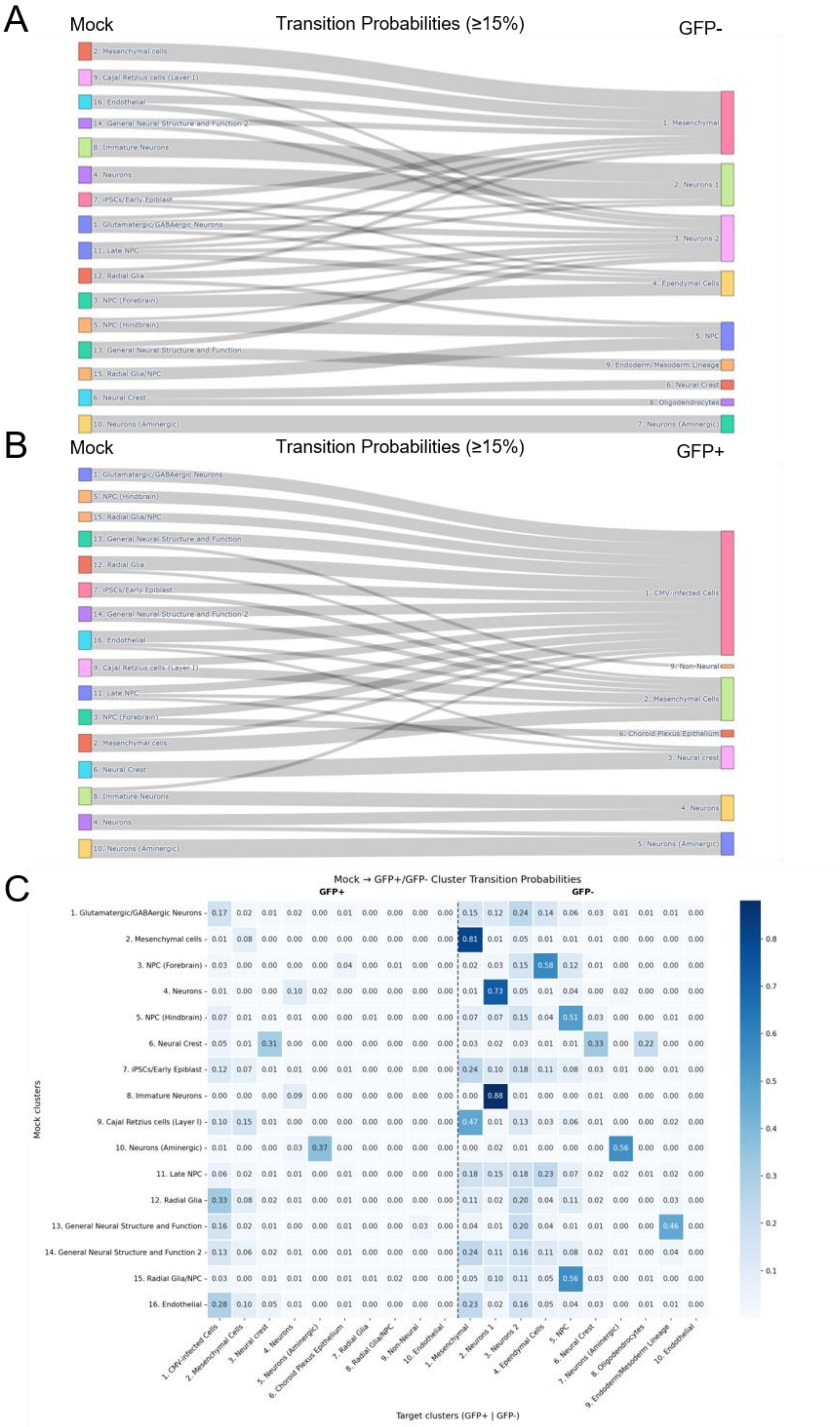
A
Mock
Transition Probabilities (≥15%)
GFP-
2. Mesenchymal cells
9. Cajal Retzius cells (Layer I)
16. Endothelial
14. General Neural Structure and Function 2
8. Immature Neurons
4. Neurons
7. iPSCs/Early Epiblast
1. Glutamatergic/GABAergic Neurons
11. Late NPC
12. Radial Glia
3. NPC (Forebrain)
5. NPC (Hindbrain)
13. General Neural Structure and Function
15. Radial Glia/NPC
6. Neural Crest
10. Neurons (Aminergic)
1. Mesenchymal
2. Neurons 1
3. Neurons 2
4. Ependymal Cells
5. NPC
9. Endoderm/Mesoderm Lineage
6. Neural Crest
8. Oligodendrocytes
7. Neurons (Aminergic)
B
Mock
Transition Probabilities (≥15%)
GFP+
1. CMV-infected Cells
9. Non-Neural
2. Mesenchymal Cells
6. Choroid Plexus Epithelium
3. Neural crest
4. Neurons
5. Neurons (Aminergic)
C
Mock → GFP+/GFP- Cluster Transition Probabilities
GFP+
GFP-
Mock clusters
Target clusters (GFP+ | GFP-)
7. Radial Glia
8. Radial Glia/NPC
10. Endothelial
10. Endothelial

**S4 Fig. Comparing MOSCOT transition probability mapping between mock to GFP-positive and mock to GFP-negative.** (A) Sankey plot showing transition probabilities between clusters in the Mock and the GFP-negative subset of HCMV-infected neural organoids. Bands represent the probability of Mock clusters transcriptionally transitioning to GFP-negative clusters following perturbation. Band width is proportional to the transition probability value shown in the transition probability matrix in Figure 3. A threshold was set at >15% transition probability for visualization of only larger bands. Between the Mock and GFP-negative secondary control, clusters tend to map onto like clusters, indicating a high degree of transcriptional similarity between those clusters. (B) Sankey plot showing transition probabilities between clusters in the Mock and the GFP-positive subset of HCMV-infected neural organoids. (C) Heatmap of transition probability matrix from a combined MOSCOT analysis of Mock to concatenated GFP-positive and GFP-negative data.

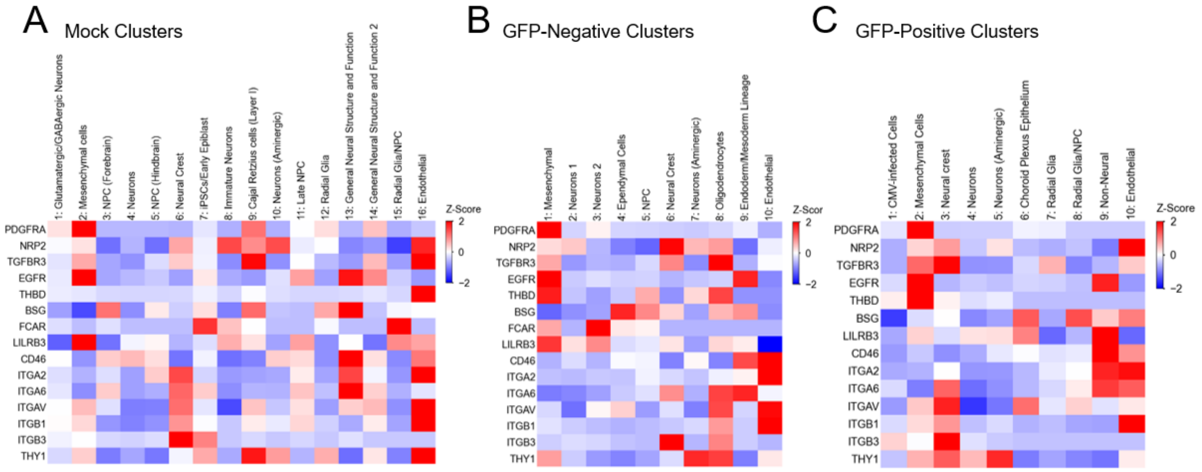


**S5 Fig. Expression of HCMV viral entry receptors across infected cell types.** Heatmaps showing the expression of various HCMV viral entry receptors in (A) mock cell type clusters, (B) GFP-negative cell type clusters, and (C) GFP-positive cell type clusters.

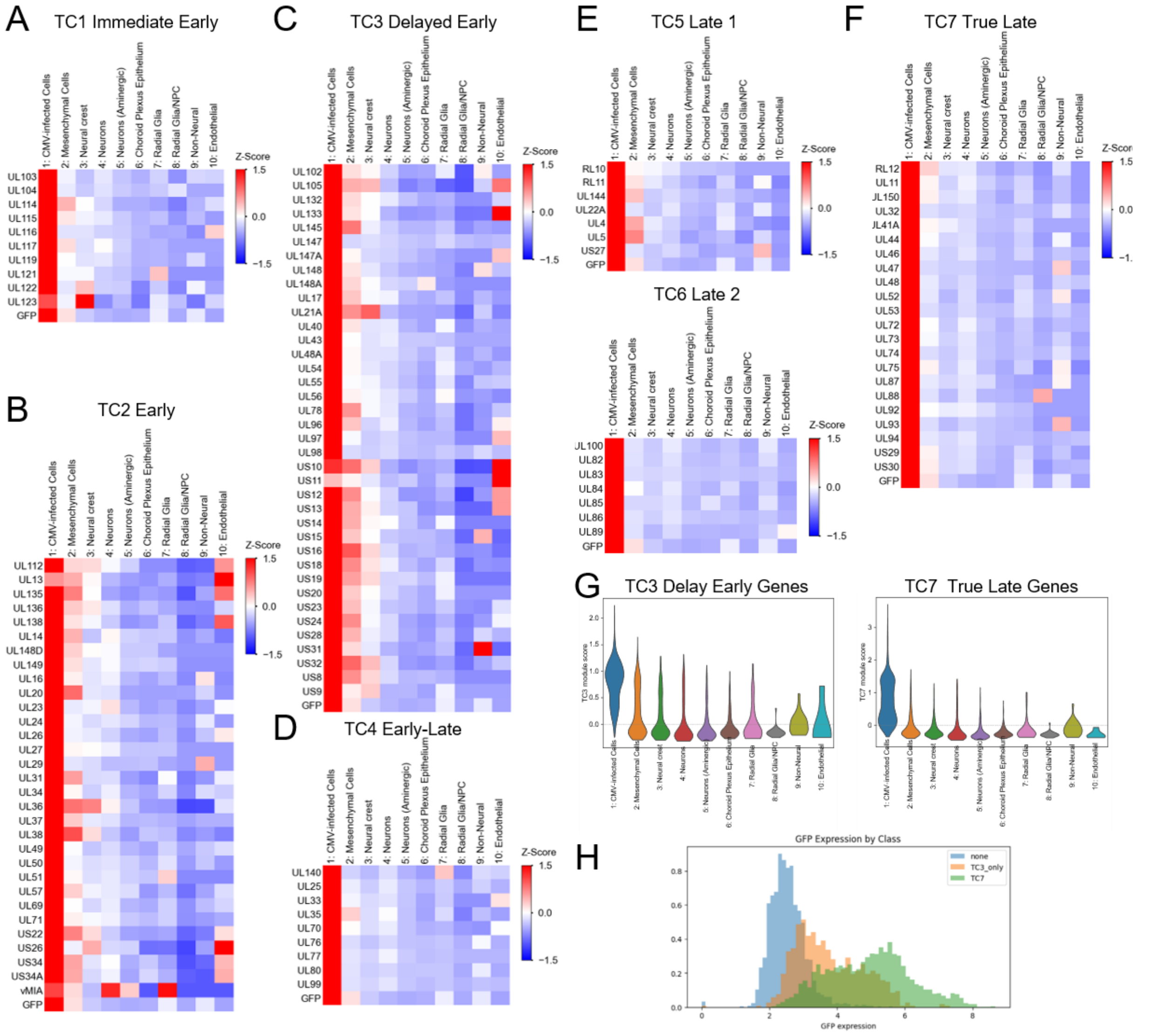


**S6 Fig. Expression of viral gene temporal classes across infected cell types.** (A-F) Heatmaps of HCMV viral gene expression grouped by temporal class (TCs). (G) Violin plots showing module scores ofHCMV viral genes comprising the TC3 and TC7 temporal classes. (H) Histogram showing GFP transcript expression in cells derived from the GFP positive subset, grouped by expression of TC7 genes, expression of TC3 genes but not TC7 genes, and expression of neither TC3 nor TC7 genes. GFP transcript levels increase as a function of TC progression.

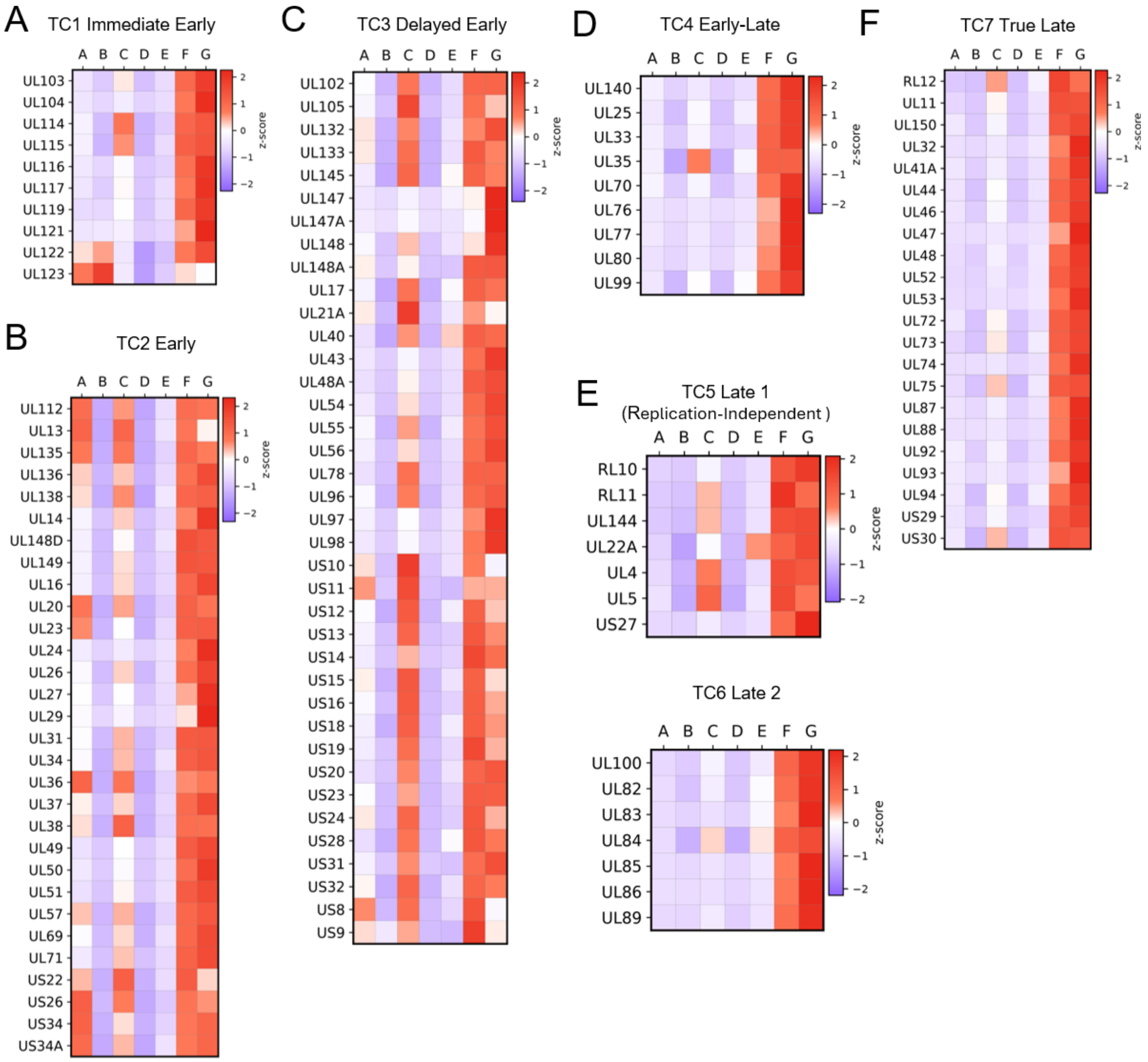


**S7 Fig. Viral gene expression across viral state clusters grouped by temporal class (TC).** (A-F) Heatmaps showing the z-scored expression of viral genes (grouped by temporal classes) across each viral state cluster. Z-scores were computed against the mean expression of a given gene across all viral state clusters.

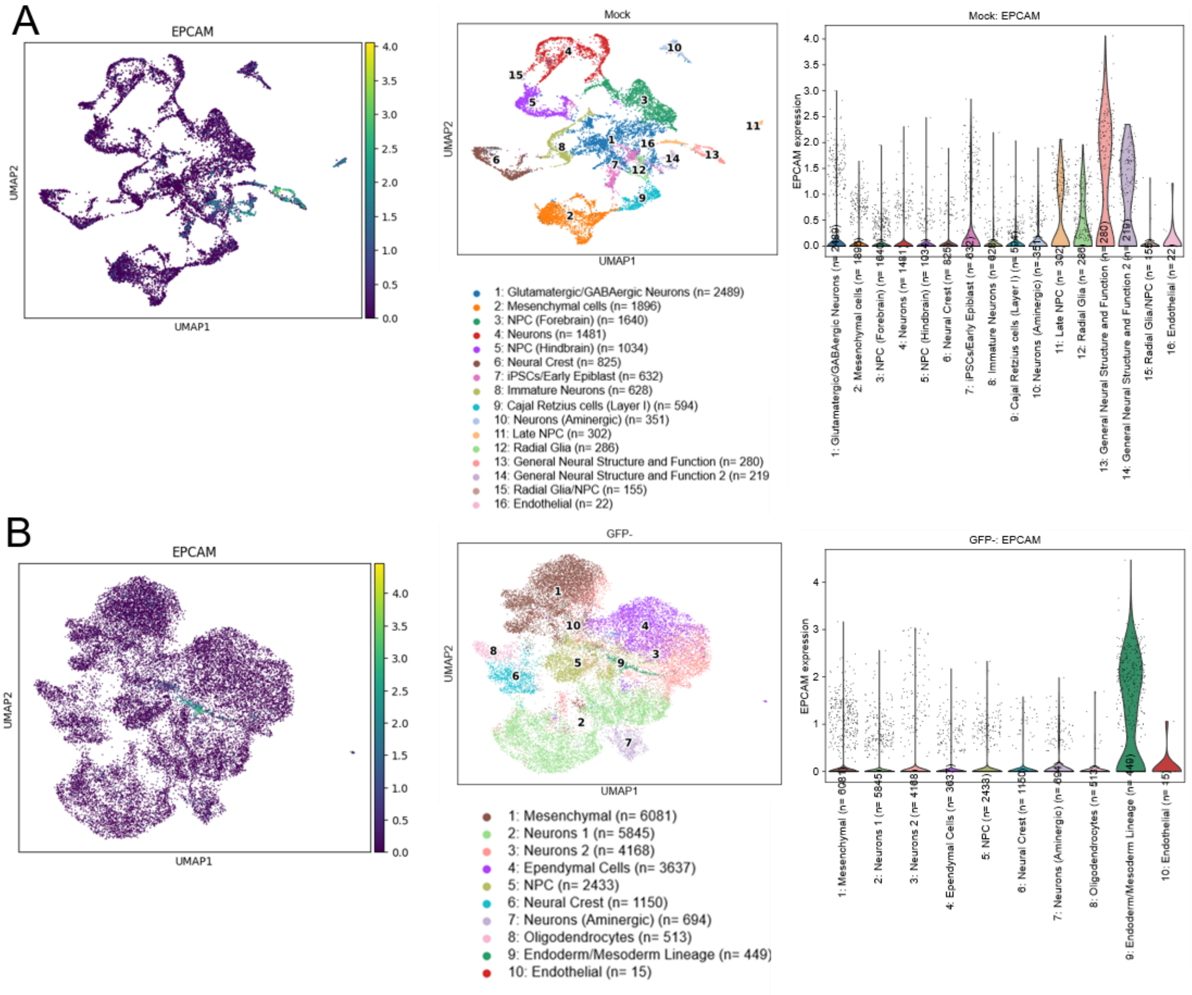


**S8 Fig. Comparison of EpCAM expression across mock and GFP-negative cortical organoids.** UMAP projection and violin plots showing EpCAM expression across different cell type clusters in (A) the mock cortical organoid and (B) GFP-negative subset of the HCMV-infected cortical organoid.
